# Structural insights into maturation and translation of a plant mitoribosome

**DOI:** 10.1101/2024.10.28.620559

**Authors:** Vasileios Skaltsogiannis, Tan-Trung Nguyen, Nicolas Corre, David Pflieger, Todd Blevins, Yaser Hashem, Philippe Giegé, Florent Waltz

## Abstract

Ribosomes are key molecular machines that translate mRNA into proteins. Mitoribosomes are specific ribosomes found in mitochondria, which have been shown to be remarkably diverse across eukaryotic lineages. In plants, they possess unique features, including additional rRNA domains stabilized by numerous pentatricopeptide repeat proteins. However, the molecular mechanisms of translation by plant mitoribosomes remain largely unknown. Here, we use cryo-electron microscopy to provide a high-resolution structural characterization of the flowering plant mitoribosome, in translating and maturation states. The structure reveals the mitoribosome bound to tRNA in the peptidyl site, along with a segment of mRNA and a nascent polypeptide. Moreover, we identify an extensive set of ribosomal RNA modifications that we confirmed by nanopore sequencing. Additionally, we observe a late assembly intermediate of the small ribosomal subunit, in complex with the RsgA assembly factor. This reveals how a plant-specific extension of RsgA blocks the mRNA channel to prevent premature mRNA association before complete small subunit maturation. Our findings elucidate key aspects of translation in flowering plant mitochondria, revealing its distinct features compared to other eukaryotic lineages.

## Introduction

Mitochondria are vital organelles found in nearly all eukaryotic organisms, where they play a central role in energy production and metabolism^1,2^. Through oxidative phosphorylation, these organelles are responsible for recycling ADP to ATP, the energy currency of the cell^3^. They are also involved in numerous core cellular functions, such as calcium signaling, apoptosis, the synthesis of heme and other essential metabolites^4^. In photosynthetic eukaryotes, such as plants and algae, the role of mitochondria is closely linked to that of chloroplasts^5,6^. While chloroplasts are responsible for producing energy-rich molecules, they are not capable of carrying out all the necessary metabolic processes on their own^7^. To overcome this limitation, photosynthetic organisms rely on their mitochondria to carry out these metabolic processes^8^.

Due to their endosymbiotic origin, mitochondria possess their own genomes and specific gene expression machineries^9^. Among these, the mitochondrial ribosome (called mitoribosome hereafter) is an essential molecular machine that synthesizes proteins within the organelle. In particular, some genes encoding subunits of respiratory complexes are encoded in mitochondrial genomes and mitoribosomes are thus essential for the biogenesis of oxidative phosphorylation complexes^10,11^. Because respiratory proteins are often intrinsic membrane proteins, mitoribosomes have evolved and specialized for the synthesis of these hydrophobic proteins^12–15^. Recent advances in cryo-electron microscopy (cryo-EM) have enabled the study of mitoribosomes at high resolution, providing valuable insights into their composition, architecture and function. Mitoribosomes were characterized in a wide diversity of eukaryotes, i.e., in mammals, fungi, kinetoplasts, ciliates, flowering plants and green alga^13,16–23^. These studies have revealed the remarkable diversity of mitoribosome composition and structures^24,25^. Still, a common feature of mitoribosomes is their high protein content, with at least 60 novel ribosomal proteins occurring in the respective mitoribosomes as compared to the bacterial ribosome^26^. Interestingly, helical repeat proteins such as pentatricopeptide repeat (PPR) proteins are prevalent among mitoribosome-specific proteins^9^. The initial description of mammalian mitoribosomes has shown that ribosomal RNA content is extremely reduced in these eukaryotes^16,17^. This led to the hypothesis that the contraction of rRNA in mitoribosomes could be compensated by increased numbers of ribosomal proteins. Likewise, this suggested that mitoribosomes could follow an evolutionary trend to reduce rRNA size, that could ultimately result in the complete loss of rRNA^27^. Even though this model applies to several eukaryotes, it was contradicted by the characterization of fungal and, in particular, of flowering plant mitoribosomes that contain both increased numbers of proteins and expanded rRNAs^18,20,28^.

Flowering plant mitoribosomes were characterized at the biochemical and structural level in the closely related Brassicaceae species Arabidopsis and cauliflower^20,29^. This revealed a very distinctive, enlarged mitoribosome with several plant-specific ribosomal proteins, among which 10 are PPR proteins. The small subunit (SSU) is particularly singular as it is larger than the ribosomal large subunit (LSU) and contains a distinctive plant-specific elongated domain on the head of the SSU^30^. Despite these advances, many features of plant mitochondrial translation remain elusive. For instance, how these ribosomes assemble, are tethered to the membrane or the way mRNAs are recruited to the SSU remain speculative. Likewise, the molecular details of the actual translation process catalyzed by the plant mitoribosome are unknown.

To address some of these open questions, we produced a sub 3 Å cryo-EM structure of the flowering plants mitoribosome. It reveals new details about rRNA modifications and previously unaccounted r-proteins. Along with the improved resolution, we also describe the structure of a translating mitoribosome with mRNA and a P-site tRNA, stalled in the presence of chloramphenicol. Moreover, a very late assembly intermediate of the SSU, bound with the assembly factor RsgA, was obtained revealing the function of its plant-specific domain and thus for the first time the mode of action of this factor in the context of a mitoribosome.

## Results & Discussion

### Structure determination

To improve our understanding of flowering plant-specific translation processes, we purified mitochondria from cauliflower and subsequently purified mitoribosomes (see methods). We purified them in the presence of chloramphenicol or non-hydrolysable GTP analog to stall translation and/or other processes that might require GTP hydrolysis. Four datasets were collected (see methods), resulting in the three main structures presented here: a full ribosome structure without tRNA resolved to 2.1 Å (Supplementary Fig 1 and 2), a stalled ribosome harboring a P-site tRNA and mRNA resolved to 3.0 Å (Supplementary Fig 3), and a late SSU assembly intermediate in presence of the assembly factor RsgA obtained at 2.2 Å (Supplementary Fig 4).

Given the large flexible extensions of this mitoribosome, we first used 3D Flexible Refinement^31^ (Movie 1), to investigate the major modes of motion of its different domains. On top of the classical ratcheting motion, we also observed the head extension motion, previously described, but also the movement of the SSU foot and of the LSU back extension. Then, extensive masking and focused refinements revealed parts of the mitoribosome that remained poorly resolved in the previous studies, notably the SSU head extension. Given the high local resolution for the different parts of the ribosome (up to 1.8 Å locally, permitting to place ions and solvent molecules (Fig. 1c and Supplementary Fig. 1 and 2)), we decided to adopt an unbiased approach to rebuild and identify the ribosomal proteins. We used ModelAngelo^32^ to sequence the proteins directly from the density and the resulting models were verified against the protein candidates previously obtained by MS^29^. This allowed the identification of a total of 85 proteins (47 in the LSU and 38 in the SSU) which we consider to be the complete set of core r-proteins of this mitoribosome (complete table of proteins Supplementary Table 2 and 3). Compared to the previous structure, we discovered new proteins and features that were previously unidentified.

**Figure 1.**
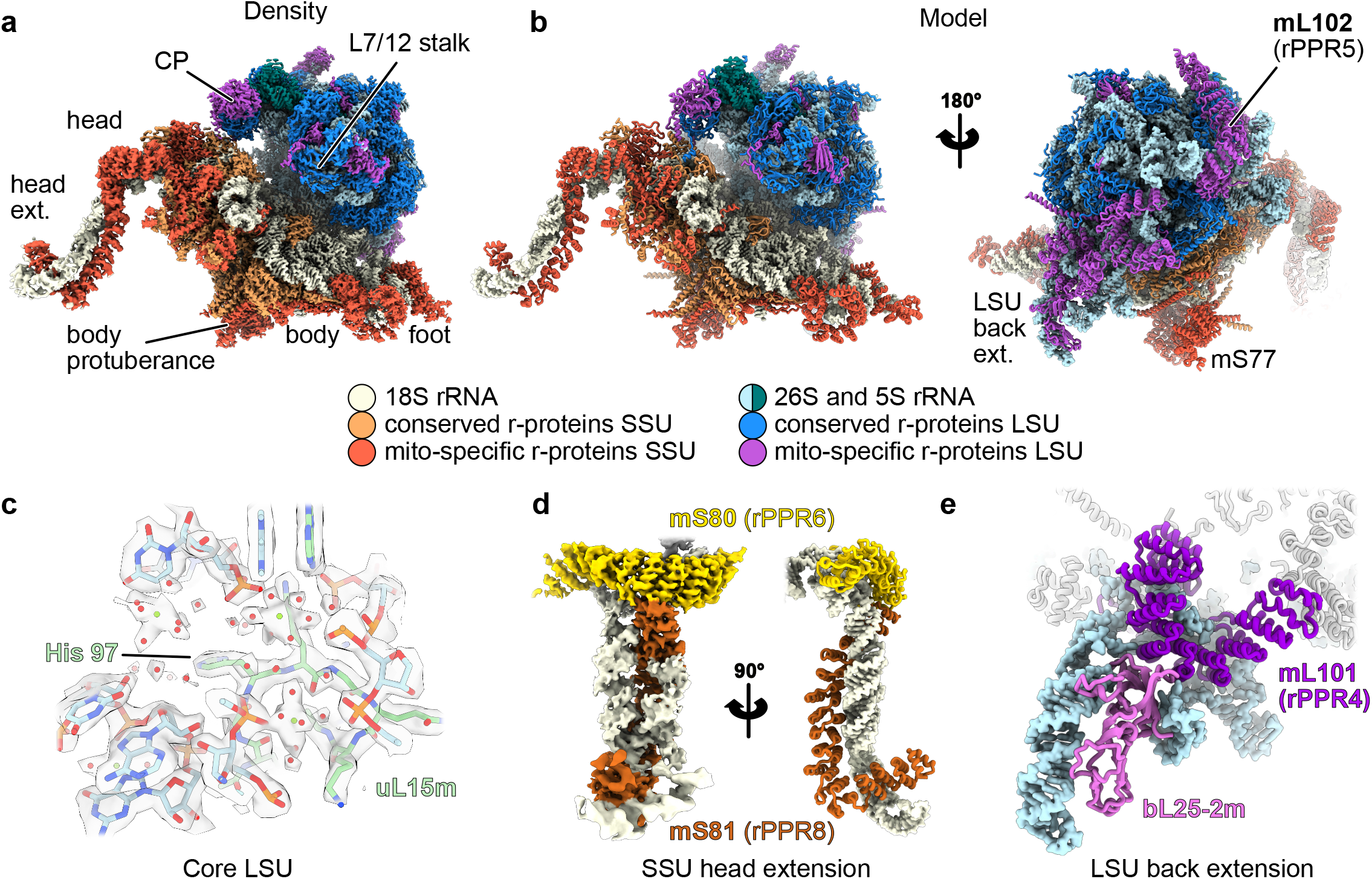
High resolution structure of the plant mitoribosome. **a** Composite high-resolution map of the cauliflower mitoribosome with components of the SSU shown in beige, coral and red and components of the LSU shown in blue shades and purple. **b** Resulting atomic model with proteins shown in cartoon representation and rRNA shown as surface. **c** Close-up view of the core of the LSU, showing solvent molecules and ions. **d** View of the SSU head extension, on the left as a density, and on the right a rotated view as an atomic model. **e** View of the LSU back extension, with the remodeled rRNA Domain III and the newly identified r-protein bL25-2m.

**Figure 2.**
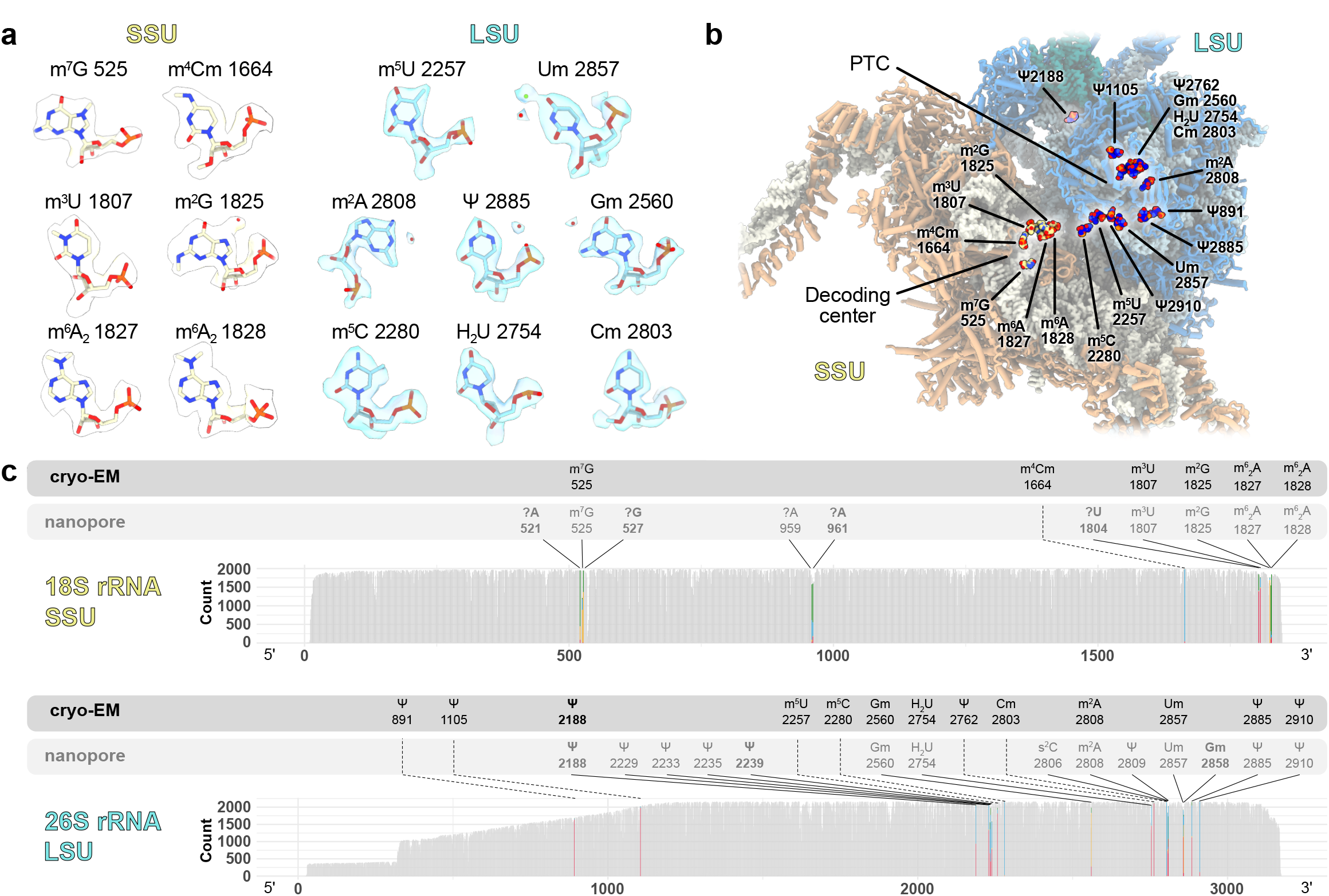
Plant mitoribosome rRNAs are extensively modified. **a** rRNA modifications identified from the cryo-EM map are shown in their respective densities with modifications in the LSU 26S rRNA shown in light blue and in the SSU 18S rRNA shown in beige. **b** Positions of the newly identified modifications are indicated on the structure of the mitoribosome. **c** Modifications identified by nanopore and cryo-EM. Plots represent the base composition of native rRNA sequences compared with in vitro rRNA transcripts (IVT) sequences obtained by nanopore direct RNA sequencing. Base positions in the rRNA sequences are shown on the x-axis, while number of read counts are shown on the y-axis. Colored positions reveal rRNA modification positions detected either by discrepancies between native and IVT sequences or by identification in the cryo-EM structure. Color code corresponds to mis-called bases as shown in Supplementary Fig. 9 and 10. The nature and positions of modifications observed in the structure are indicated in black, while modifications observed by nanopore are indicated in grey.”?” correspond to novel RNA modifications that could not be characterized. The 7 modifications that are specific to the plant mitoribosome are indicated in bold. Details for each position identified by nanopore sequencing are shown Supplementary Fig. 9 and 10 and listed in Supplementary Table 4.

### Newly resolved features

In the large subunit, the remodeled domain III of the 26S rRNA on the LSU back is highly flexible (see Movie 1) and was thus poorly resolved. There, the rRNA is remodeled mainly by PPR proteins mL104 (rPPR9) and mL101 (rPPR4). Using focused refinement, we resolved it in greater detail, notably their interaction with the rRNA. Additionally, it also revealed the presence of a new r-protein (Fig. 1e and Supplementary Fig. 5a-d). This protein is an isoform of bL25m, that we name bL25m-2, interacting with both the remodeled domain III and mL101. There, it shapes and stabilizes the remodeled domain III by mimicking the interaction of bL25m with the 5S and 26S rRNA located between the L7/L12 stalk and the base of the Central Protuberance (CP) (Supplementary Fig. 5d). Focused refinement on the L7/L12 stalk revealed the presence of mL54 (conserved with algae, mammals and yeast) which was previously identified only by mass spectrometry (Supplementary Fig. 5e). uL2m is split into two parts in the flowering plants, one encoded in the mitochondrial genome (N-terminal portion) and one in the nuclear genome (C-terminal portion), and forms an intricate network of protein-protein interactions with multiple r-proteins. First it interacts with mL86, and then extends far away from the uL2 region to closely interact with bL28m, bL9m as well as mL102 (Supplementary Fig. 6c).

The largest PPR protein of the LSU, mL102 (rPPR5), contacts the 26S rRNA at different positions of the LSU plus several proteins, namely uL15m, mL59/mL64 and the N-ter portion of uL2m. mL102 contributes to the reorganization of the domain I of the 26S rRNA mostly through its N-ter part (Supplementary Fig. 6a). We could resolve how the domain I plant-specific helices p15, p17 and p21-22 are folded. This portion of the rRNA and the protein mL102 are less well resolved because they extend far into the solvent, but also because they might be involved in membrane interaction during translation^20^. In the stalled state of the mitoribosome (described later) we could also partially resolve uL1m on the L1 stalk (Supplementary Fig. 6b).

The small subunit is where the model improved the most, allowing clear identification of several proteins that were previously modeled but not identified. The most drastic improvement can be observed on the SSU head extension (Fig. 1d). In flowering plants, the 18S rRNA harbors a large extension at the tip of the SSU’s head, rooting from h39 of domain 3’M. Here we resolve this specific domain to approximate 4Å (compared to the previously reported 8Å). This allowed, both thanks to improved resolution and improved structure prediction using AlphaFold2^33^, the unambiguous identification of the two r-proteins located there (rPPR6 and rPPR8) (Supplementary Fig. 7a), as well as building a more accurate rRNA. mS80 (rPPR6) is positioned at the base of the head extension, whereas mS81 (rPPR8) wraps around and follows the shape of the head extension. Two PPR modules are found, the longest one (aa 219-656) follows the two parallel RNA helices, while the second one (aa 76-218) is bent almost 90° relative to the first one, and helps form the tip of the SSU head extension.

On the opposite side of the SSU, at the foot, we clearly identify mS76 (rPPR1) and mS83 (rPPR10), which were previously modeled as Unknown proteins (Supplementary Fig. 7b). mS76 contributes to the stabilization of the h6 helix of domain 5’ of the 18S rRNA whereas mS83 mostly contributes to the stabilization of h44 tip, as well as h6. A recent study has performed the functional analysis of mS83 (called mTRAN by authors)^34^. A global impairment of mitochondrial translation in mS83 mutants was observed and mS83 was found to be able to bind RNA *in vitro*. The authors proposed that mS83 is a general factor for translation initiation in plant mitochondria. However, here we place mS83 at the tip of the foot of the SSU, interacting with plant specific rRNA expansion segments, which is not consistent with a role in translation initiation.

At the exit of the mRNA channel we find mS77, another PPR protein (Supplementary Fig. 7c). It wraps around the SSU body, rooting from the SSU protuberance, where it is anchored and extends in the back of the SSU, close to the mRNA exit channel where its PPR domain is found. Originally listed as two proteins in UNIPROT (Q8LEZ4, NFD5, N-ter part, making part of the body protuberance and Q940Z1, PPR51, C-terminal part containing the PPR motifs), we instead describe it as a single protein like in the UNIPROT entry A0A654ECT9. Moreover, at low threshold we also observe an additional density at the tip of the PPR domain, suggesting that this protein might act as a platform, to recruit specific plant translation factors, or allow polysome formation (Supplementary Fig. 7c). Next to this protein, we also identify a small portion of the N-terminal part of mS86 (GRBP6), (residues 1 to 28, the protein being 155aa), sandwiched in between uS2m and mS23 (Supplementary Fig. 7c). mS86 was previously identified through mass spectrometry but not resolved in the structure. Its position next to the mRNA exit channel may suggest an interaction with mRNA, like mS77, or a role in polysome formation. These additional proteins might also have been recruited to the exit channel to compensate for the absence of uS1.

Additionally, on the head of the SSU, we fully resolve the composition of the protein protuberance located close to the beak. It is formed by mS31/mS46, that is also newly identified, mS35, and uS3m, which was predicted to form part of this protuberance (Supplementary Fig. 8b). Together they form an intricate network of proteins close to the mRNA entrance channel. Finally, the improved resolution reveals the contacts between the LSU’s CP to SSU head via bL31m, mL40 and mL46, and unravel the presence of ATP in mS29 (Supplementary Fig 8a), that is also present in human mitoribosomes^35^.

Altogether, out of the 85 mitoribosomal proteins, 53 are conserved with bacteria, 22 are shared with mitoribosome of other eukaryotic species and 10 proteins are specific to flowering plants, among which 8 are PPR proteins (Supplementary Table 2 and 3).

### rRNA modifications of the plant mitoribosome

rRNA modifications are hallmarks of subunit maturation towards a translationally active ribosome^36,37^. To date, the rRNA modifications present in plant mitochondria have not been described. Our map resolution allowed us to identify in total 19 rRNA modifications, 6 in SSU and 13 in LSU (Fig. 2a-b). Additionally, we used nanopore direct RNA sequencing of the native cauliflower rRNAs to confirm the modifications detected in the structure or identify others when the map resolution was not adequate for their visual identification. Nanopore RNA sequencing has been successfully applied before to detect RNA modifications, for instance in ribosomes^38–41^. This second approach indicated the presence of 24 rRNA modifications, 10 in SSU and 14 in LSU (Fig. 2c). In total, structure inspection and nanopore sequencing resulted in the identification of 31 rRNA modifications, 11 being in the SSU and 20 in the LSU, as presented in Figure 2. Surprisingly, this number deviates considerably from the ones of the scarcely modified mammalian or yeast mitoribosomes that contain 10 and 5 rRNA modifications, respectively^39,42–45^.

We found 9 types of base modifications, including an extensive network of pseudouridines (Ψ) and 3 types of 2’-O ribose methylation (Nm). Similar to their bacterial and cytosolic counterparts, rRNA modifications of the plant mitoribosome are located in clusters close to functionally important sites (Fig. 2b). Near the decoding center, we located the universally conserved methylated residues m^6^_2_A1827 and m^6^_2_A1828. These adjacent modified adenosines aid the P-site in acquiring its mature conformation by establishing contact between helices h44 and h45. In this vicinity, m^2^G1825 of h45 and two modifications located at the base of h44, m^3^U1807 and m^4^Cm1664 were further identified, with the former two being characterized for the first time in a mitoribosome (Fig. 3d). m^4^Cm1664 interacts directly with the mRNA and has been proposed to fine-tune the decoding center in bacteria by preventing translation initiation at the alternative initiation codon AUU and by maintaining the reading frame^46^.

**Figure 3.**
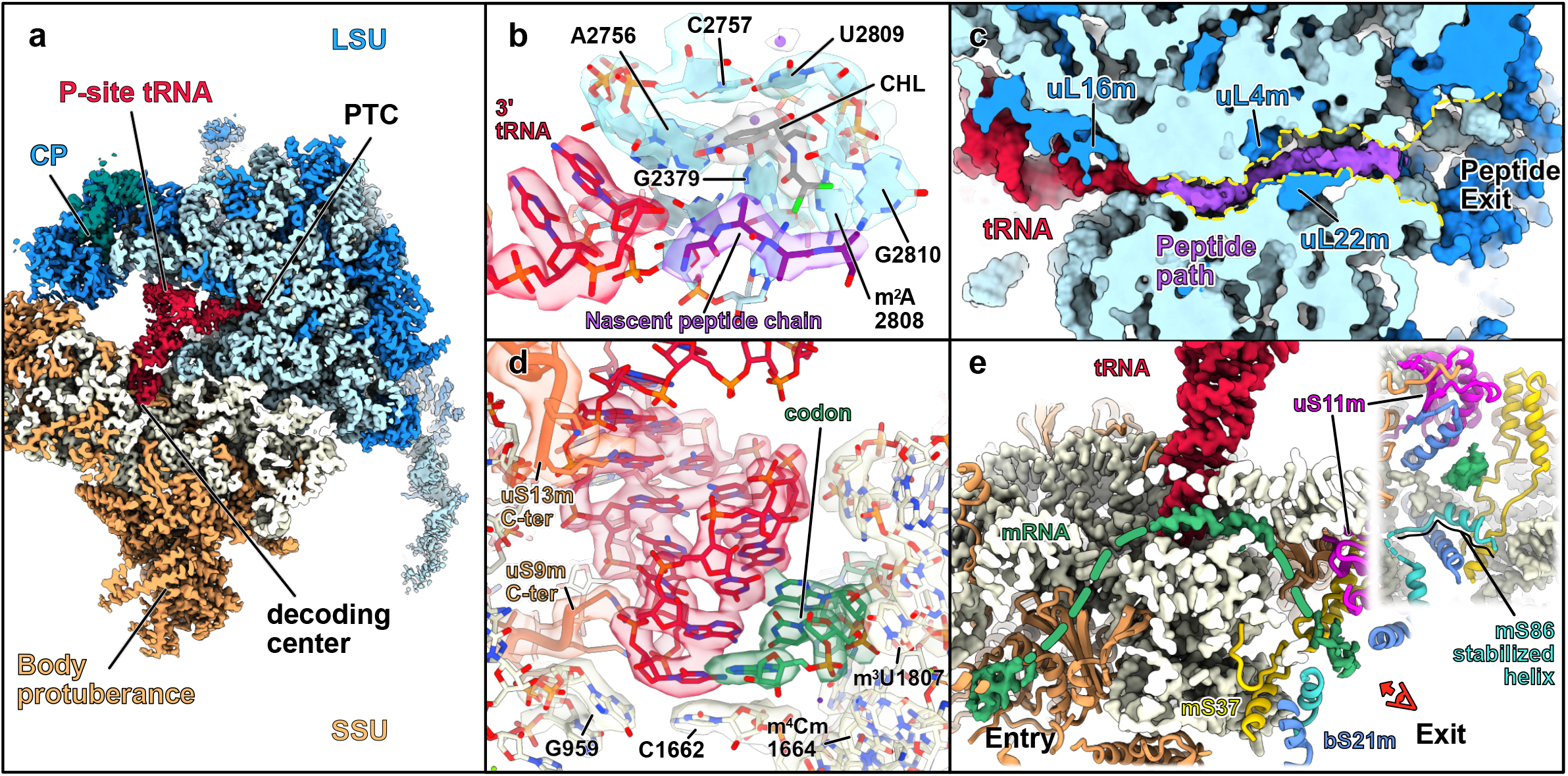
Plant mitoribosome in a translating state. **a** Composite map of the mitoribosome with the SSU shown in beige, the LSU in blue and a tRNA in the P-site shown in red. **b** Close-up view of the PTC with bound chloramphenicol (CHL) shown in grey and the stalled nascent peptide chain in purple, shown in their density. The nucleotides forming the chloramphenicol binding pocket are annotated and shown in their density. **c** Cut view of LSU, showing the peptide path, with the nascent peptide chain in purple. **d** Close-up view of the decoding center highlighting the interaction between the codon in green and the anticodon in red as well as uS13m and uS9m probing the anticodon loop, shown in their density. **e** Cross-section view of the SSU highlighting the mRNA channel, with resolved mRNA residues depicted in green at the decoding center, as well as at the channel’s entry and exit points. The inset provides a close-up view of the mRNA channel, viewed from the red eye, illustrating the stabilization of the mS86 helix in the presence of mRNA.

Another cluster of rRNA modifications was observed around the peptidyl transferase center (PTC). Among them, we were able to identify the highly conserved Gm2560 and Um2857 at the helices 80 and h92, respectively. Gm2560 is in the P-loop region and supports the positioning of the P-tRNA by interacting with its CCA-end, whereas Um2857 resides in the A-loop region and assists in the accommodation of the tRNA at the A-site. Interestingly, mammalian mitoribosomes possess an additional 2’-O methylation at the adjacent G2858 residue, which we could not model due to lower resolution in that region. Nonetheless, our nanopore data point to the existence of a modification on this specific residue, thus we do not rule out the existence of Gm2858. The A-tRNA accommodation is further stabilized by the pseudouridine Ψ2885, similar to human mitoribosomes^45^. The domain IV of the LSU rRNA establishes the intersubunit bridges of group B2, crucial for ribosome assembly and translation. In this region, we identified the methylated sites m^5^U2257 (h70) and m^5^C2280 (h71) as well as the pseudouridine Ψ2188 (h68) which is plant-specific. Moreover, our nanopore data suggests the potential presence of four additional pseudouridylation sites in helix 69, although these sites could not be confirmed by cryo-EM, as the resolution is lower in that area of the map. Finally, the presence of the modified nucleotide m^2^A2808 was observed at the PTC (Fig. 3b). The methyl group of m^2^A2808 extends its stacking range allowing the interaction with A2377, thus promoting the folding of their cognate single-stranded rRNA segments to form the peptide exit tunnel wall.

Overall, the plant mitoribosome is highly modified and has retained a significant number of the modifications present in its bacterial ancestor. Thus, it appears to have followed an evolutionary route that allowed for the preservation of many bacterial modifications in direct contrast to its mammalian and yeast counterparts who were able to retain only a few. This also suggests that many bacterial modification enzymes must be conserved in plants, or have been replaced by new ones, to maintain these modifications.

### Translating ribosome

To trap ribosomes in translation states, we collected a dataset for mitoribosomes purified from mitochondria and ribosomes treated with chloramphenicol, a classical inhibitor of bacterial-type ribosomes. Chloramphenicol binds to the PTC of the ribosome, stalling translation^47^. During data processing, we identified a subclass corresponding to mitoribosomes in a P-site tRNA-accommodated state, which we resolved to approximately 3 Å resolution (Fig. 3 and Supplementary Fig. 3). This allowed us to build a detailed model that includes the tRNA, the messenger RNA path, the binding site of chloramphenicol, and the peptide route through the peptide channel.

At the PTC, within the LSU, we fully resolved the binding site of chloramphenicol, responsible for stalling the ribosome in the P-site tRNA-accommodated state (Fig. 3b). At this site, the first four amino acids of the nascent peptide chain were clearly visible. Since this represents a mix of averaged densities from different nascent proteins, we modeled it as poly-alanine. At lower threshold, we could observe the complete peptide chain extending from the PTC to the peptide exit via the peptide tunnel. Interestingly, the structure of the peptide tunnel and exit was found to be very similar to that of bacterial ribosomes, with less extreme adaptations compared to animals, yeast, and green algae^13,28,48^. This may be related to the fact that flowering plants, like bacteria, also translate mRNA encoding soluble proteins, unlike some of the organisms mentioned previously^30^.

Since mitoribosomes were treated with chloramphenicol, we were able to resolve the antibiotic’s binding site, revealing that its mode of action is nearly identical to its binding in classical bacterial ribosomes. This is not surprising, as the PTC is one of the most highly conserved regions of the ribosome, alongside the decoding center^49^. The chloramphenicol binding in the mitoribosome is positioned similarly to its location in bacterial ribosomes, reinforcing the conserved nature of its inhibitory mechanism across species.

In the decoding center of the small subunit (SSU), we also observed the mRNA being translated, from which we resolved six nucleotides, three of which were interacting with the anticodon loop of the stalled tRNA (Fig. 3d). Similar to the peptide observed at the PTC, these nucleotides represent an averaged mix of densities from different mRNAs. Nonetheless, we were able to observe how they are accommodated within the small subunit. Here, conserved residues of the 18S rRNA, G956, and C1746 help stack and position the tRNA. The ribosomal protein uS13m probes the base of the tRNA anticodon region with its C-terminal extension, behaving in a manner identical to bacterial ribosomes, while uS9m, which has a slightly extended C-terminus compared to its bacterial counterpart, probes a different region of the tRNA anticodon (Fig. 3d).

At a lower threshold, we could also visualize densities corresponding to mRNA at the entry and exit sites of the mRNA channel, allowing us to trace the full putative mRNA path within the SSU (Fig. 3e). At the entry point, the conserved core of proteins uS4m, uS5m, and uS3m (located on the head of the SSU) form the majority of the entry channel. Extensions of uS5m and a portion of uS4m, which form parts of the SSU body protuberance, further delineate and extend the entrance of the mRNA channel. On the SSU head, uS3m, together with mS31/46 and mS35, contributes to the SSU head protuberance near the beak. Notably, 300 amino acids from the N-terminus of mS31/46 remain unresolved and point toward the solvent, in close proximity to the mRNA entrance (Supplementary Fig. 8b). Altogether, the SSU body and head protuberances located near the mRNA entry channel could play a role in species-specific recruitment of mRNA or polysome formation, which has been observed *in situ* in Chlamydomonas^50^.

On the opposite side, at the mRNA exit, the ribosomal proteins uS11m, bS21m, and the mitochondria-specific mS37 form the exit channel. Additionally, we observed a larger portion of mS86 in the ribosome charged with tRNA and mRNA compared to empty ribosomes. An additional helix in mS86 is stabilized up to residue 54 (compared to 28 amino acids in the empty ribosome) at the mRNA exit channel (Fig. 3e). The fact that this protein is stabilized in the stalled ribosomes, where mRNA is present, supports the hypothesis that mS86 may be important for mRNA interaction and/or polysome formation.

### A late SSU assembly intermediate

Mitoribosome biogenesis is coordinated by the orchestrated action of a series of assembly and auxiliary factors that guide the rRNA to adopt its mature conformation and in parallel drive the integration of the complete set of ribosomal proteins to generate two functional ribosomal subunits. Structural and mechanistic insights into the assembly pathway of mitoribosomes have been recently provided in kinetoplastid^22,51–54^, human^36,37,55–62^ and yeast^36,63^. Nevertheless, what assembly factors participate in the biogenesis of plant mitoribosomes and the mechanisms under which they operate, remain elusive. To address these questions, we tried to capture native assembly intermediates by stalling the assembly process of the plant mitoribosome. We targeted assembly factors of the GTPase family, known to be important enzymes for ribosome maturation^64^. To do so, we isolated mitoribosomes in the presence of a non-hydrolysable GTP analog, GTPγS. Even though no clear new state of the fully assembled mitoribosome could be identified, analysis of the GTP analog SSU only fraction revealed a subclass of 208,855 particles with an additional density located near the decoding center (Fig. 4a). This class reached 2.2 Å resolution globally, allowing us to directly identify the additional factor from the density as the assembly factor RsgA. This is the first description of RsgA as part of an assembly intermediate in any mitoribosome. RsgA adopts a conformation where its central GTPase domain docks at the decoding center of the SSU (Fig. 4g) while its oligonucleotide/oligosaccharide binding-fold domain (OB-domain) spans towards the uS12m (Fig. 4d) and the Zinc-binding domain interacts with the head of the SSU via helix 29 (Fig. 4e). This conformation is similar to the one observed in bacterial ribosomes bound to RsgA^65–67^. The bacterial RsgA is considered to act as a late checkpoint for the maturation state of the decoding center, via its direct interaction with h44^65^. Consistent with this notion, we observed a similar interaction of Arg^84^ with A1802 of h44. Also, considering the overall maturation state of the SSU, the plant RsgA acts upon the late steps of the SSU biogenesis. Despite the similarities shared by the two homologs, plant RsgA possesses a 66 amino-acid long C-terminal extension that appears to have a dual function (Fig. 4c). First, it reinforces the positioning of RsgA by interacting with mS33 (Fig. 4d), and second adopts a certain conformation that allows its entry into the mRNA channel. Remarkably, the plant-specific extension folds into an alpha-helix that threads into the mRNA channel via its placement in between uS3m, uS5m, and uS4m (Fig. 4c). Therefore, RsgA blocks the mRNA channel and prevents the premature accommodation of the mRNA, during the assembly process. The function of sequestering the immature SSU particles from those that are fully functional is pivotal and is served by various mechanisms along the assembly pathway, mainly by making the mRNA channel inaccessible. Interestingly, during the late stages of SSU biogenesis of the human mitoribosome, this role is performed by the assembly factor RBFA, which has acquired a mitochondria-specific C-terminal extension that spans across the mRNA channel preventing the mRNA recruitment. It is apparent that the plant RsgA and the human RBFA are two evolutionary unrelated assembly factors, that have independently acquired lineage-specific extensions in their amino acid sequence. These extensions perform a novel similar function, expanding their basic functional repertoire in the assembly process. Thus, these assembly factors constitute a fine example of convergent evolution between plants and human mitochondria.

**Figure 4.**
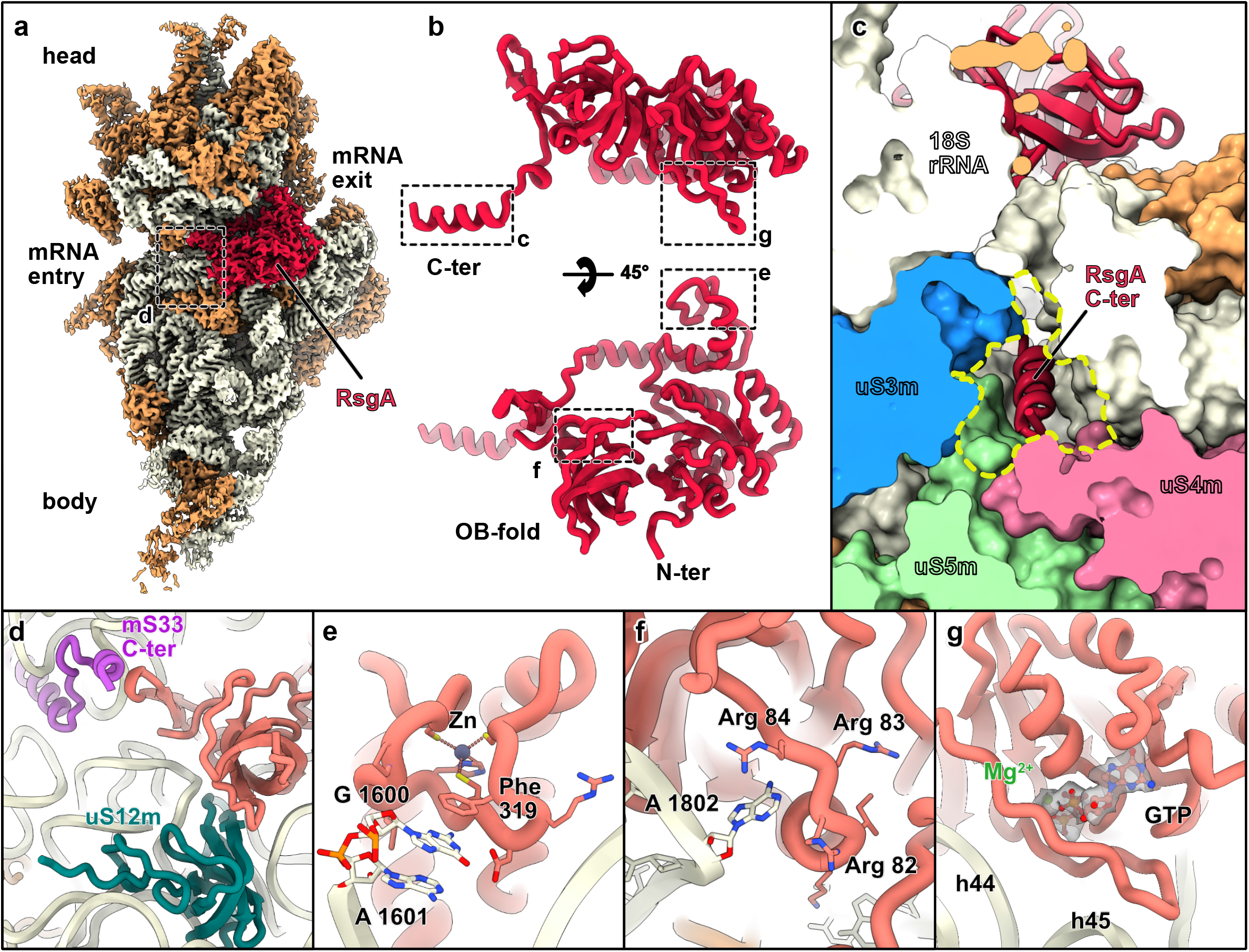
Late stage SSU maturation complex in presence of RsgA. **a** Composite map of the SSU shown in beige, with r-proteins shown in coral, rRNA shown in beige, and RsgA shown in red. **b** Close-up view of the RsgA atomic model, shown from two different point of views. Dashed boxes indicate further detailed views. **c** View of the RsgA C-terminus blocking the mRNA channel entry. **d-g** Close-up views of the different positions probed by RsgA. **d** RsgA protein-protein contacts with mS33 C-terminus and uS12m. **e** Zinc-binding domain area, with Phe319 probing G 1600 on the head of the SSU. **f** Arg 84 from the OB-fold domain probing A1802 at the base of h44. **g** GTP binding site, with GTP analog shown in its density. In that region, RsgA probes the base of h44 and the tip of h45.

## Conclusions

In this study, we provide new insights into the function and assembly of the translation apparatus in flowering plants using cryo-EM. While it was already known that plant mitoribosomes exhibit significant structural complexity, with expanded rRNA domains and numerous plant-specific ribosomal proteins, the data presented here provide significantly improved resolution and identify previously uncharacterized r-proteins. Furthermore, we detected novel features such as nucleotide modifications, revealing a high number of rRNA modifications in plant mitoribosomes as compared to mammalian and yeast mitoribosomes. These modifications are located near functionally critical regions such as the decoding center and the peptidyl transferase center, highlighting the evolutionary conservation of certain bacterial features in plant mitoribosomes. Although some modifications were acquired during the evolution of eukaryotes, most rRNA modifications represent ancestral features of mitoribosomes. Furthermore, our study identifies the plant specificities of key functional domains of the mitoribosome, in particular the decoding center and mRNA path as well as the peptidyl transfer center, further elucidating unique aspects of translation in plant mitochondria. Finally, we captured a late-stage assembly intermediate of the mitoribosome, bound to the assembly factor RsgA. This interaction reveals the mechanism by which RsgA ensures the proper maturation of the small subunit by blocking the mRNA channel during assembly, thanks to a plant-specific domain.

Altogether, this study advances our understanding of the plant mitochondrial translation apparatus structure and function, revealing novel features that distinguish plant mitoribosomes from their counterparts in other eukaryotes as well as in prokaryotes. These results show that while plant mitoribosomes have followed a distinct evolutionary path, they have preserved bacterial-like rRNA modifications in their comparatively well-conserved core domains while also incorporating unique plant-specific proteins and evolving specific assembly mechanisms. These structural features underscore the functional complexity and evolutionary adaptations of plant mitoribosomes, possibly required to fulfil plant-specific functions, such as decoding a set of mRNAs that are more diverse and more complex than in other eukaryotes.

## Supplementary Figure Legends

**<u>Supplementary Figure 1 :</u>** Single-particle data processing workflow of the high resolution mitoribosome

Graphical summary of the processing workflow described in Methods. **a** Post-processing, 2D and 3D classification. **b** Focused refinements for the LSU. For the final reconstruction, GSFSC curves are plotted, with resolution calculated at the 0.143 threshold. Local resolution plotted on the maps were generated using the built-in cryoSPARC tool with default parameters (FSC=0.5 threshold), all using the same resolution scale, with maps also shown in cut view.

**<u>Supplementary Figure 2 :</u>** FSC plots of the SSU from the high resolution mitoribosome

Focused refinements for the SSU. For the final reconstruction, GSFSC curves are plotted, with resolution calculated at the 0.143 threshold. Local resolution plotted on the maps were generated using the built-in cryoSPARC tool with default parameters (FSC=0.5 threshold), with a resolution scale different for the SSU head extension, with maps also shown in cut view.

**<u>Supplementary Figure 3 :</u>** Single-particle data processing workflow of the stalled tRNA mitoribosome

Graphical summary of the processing workflow described in Methods. **a** Post-processing, 2D and 3D classification. **b** Focused refinements for the tRNA class. For the final reconstruction, GSFSC curves are plotted, with resolution calculated at the 0.143 threshold. Local resolution plotted on the maps were generated using the built-in cryoSPARC tool with default parameters (FSC=0.5 threshold), all using the same resolution scale, with maps also shown in cut view.

**<u>Supplementary Figure 4 :</u>** Single-particle data processing workflow of the SSU with RsgA

Graphical summary of the processing workflow described in Methods. **a** Post-processing, 2D and 3D classification. **b** Focused refinements for the RsgA class. For the final reconstruction, GSFSC curves are plotted, with resolution calculated at the 0.143 threshold. Local resolution plotted on the maps were generated using the built-in cryoSPARC tool with default parameters (FSC=0.5 threshold), all using the same resolution scale, with maps also shown in cut view.

**<u>Supplementary Figure 5 :</u>** Newly resolved r-proteins in the LSU

**a-d** Newly identified r-protein bL25-2m in the LSU back extension, remodeling the 26S domain III. **a** show a close-up of bL25-2m, shown in **b**, with the experimental density. **c** Close-up view of the canonical bL25m r-protein in the mitoribosome. **d** Overlay of the remodeled domain III from **b**, shown in yellow, with the 5S and 26S rRNA from **c** showing the structural mimicry of the rRNA, with the base of p56 mimicking H4 of the 5S and p56’s tip mimicking H38, as well as p58 mimicking the base of the L7/17 stalk. **e** View of the L7/12 stalk with the experimental density showing mL54 which was previously unresolved. The different areas of the LSU are shown in the middle of the figure, with panel **b, c** and **e** positions indicated.

**<u>Supplementary Figure 6 :</u>** Improved model in the LSU

**a** Close-up view of the mL102 area. The rPPR protein mL102 interacts with three r-proteins, uL15m, mL59/64 and uL2m (N-ter mito). The N-terminal part of the protein largely interacts with rRNA and contributes to the remodeling of domain I, forming p17, p17 and p21-22, specific helices of the plant mitoribosome. Unmodeled densities of the rRNA are shown are blue dashed lines. **b** View of the L1 stalk at low threshold, revealing the presence of the uL1m protein. **c** In the plant mitoribosome, uL2m is split into two parts. What would correspond to the N-terminal part of the canonical uL2 is encoded as uL2m in the mitochondrial genome (yellow), whereas the C-terminal part is encoded in the nuclear genome (orange). The uL2m N-ter mito possesses large insertions that extend away from the uL2 core to make intricate interactions with bL28m, bL9m and mL102. Close to the uL2 core it also makes direct interactions with mL87. The different areas of the LSU are shown on the right hand-side of the figure, with panel **a, b** and **c** positions indicated.

**<u>Supplementary Figure 7 :</u>** PPR proteins of the SSU

**a** On the SSU head extension, two PPR proteins are found, mS80 and mS81. Portions of the atomic models (highlighted by dashed boxes) are shown in the experimental density. **b** At the foot of the SSU, two PPR proteins are found, m76 and mS83. Portions of the atomic models (highlighted by dashed boxes) are shown in the experimental density. **c** The newly identified r-protein mS77 (red) spans from the SSU body protuberance to the exit mRNA channel. It is composed of two main domains. Forming part of the SSU body protuberance, the N-terminal part interacts with extensions of uS5m and uS4m as well as mS47. Through a flexible portion of the protein (represented by dashed-lines) it wraps around the SSU body and its C-terminal part is composed of a long stretch of PPR repeats, interacting with mS23 and an extension of uS9m. At the mRNA exit channel a small portion of mS86 is also visible, sandwiched between mS23 and uS2m. At lower threshold, we also observe additional densities at the C-terminus of mS77, that could indicate that it serves as a binding platform for additional factors. C- and N-termini of the proteins of interest are indicated by colored N and C. The different areas of the SSU are shown on the right hand-side of the figure, with panel **a, b** and **c** positions indicated.

**<u>Supplementary Figure 8 :</u>** Improved model in the SSU

**a** Close-up view of mS29 on the SSU head, showing the ATP with Mg^2+^ in its density. **b** View of the SSU head protuberance, or S3 area. This domain is formed by a large insertion in uS3m, forming intricate interactions with the N-terminal part of mS35 and a large portion of mS31/mS46. with C- and N-termini of the proteins of interest are indicated by colored N and C. The different areas of the SSU are shown on the right hand-side of the figure, with panel **a** and **b** positions indicated.

**<u>Supplementary Figure 9 :</u>** Nucleotide frequency at positions with detected modifications

Base-called nucleotides and their percentage presented for all the positions with detected modifications of 18S rRNA (**a**) and 26S rRNA (**b**). Comparison of IVT and Native rRNAs is presented, with the latter displaying base-calling errors accompanying the presence of rRNA modifications. Positions with identified modifications from the cryo-EM map are also included. Base-called nucleotides are colored.

**<u>Supplementary Figure 10</u> :** Current intensity and nucleotide frequency at modified positions

Per-read current intensity analysis and nucleotide frequency in the 15-mer regions surrounding the modified sites of 18S rRNA (**a**) and 26S rRNA (**b**). Green lines correspond to native reads whereas brown lines to IVT reads. Thick green and brown lines represent the mean current intensity of native and IVT reads, respectively.

## Supplementary Table Legends

**<u>Table 1 :</u>** Cryo-EM validation

**<u>Table 2 :</u>** List of the LSU r-proteins

List of the r-proteins found in the LSU. Proteins are colored by conservation with the bacterial ribosome (blue) other mitoribosomes (yellow) or specific to the plant mitoribosome (red). Protein modeled in the density are indicated.

**<u>Table 3 :</u>** List of the SSU r-proteins

List of the r-proteins found in the SSU. Proteins are colored by conservation with the bacterial ribosome (blue) other mitoribosomes (yellow) or specific to the plant mitoribosome (red). Protein modeled in the density are indicated.

**<u>Table 4 :</u>** List of the rRNA modifications

Summary of the rRNA modifications detected in B. oleracea mitoribosome. B. oleracea contains 11 modified residues in the SSU and 20 in the LSU. The method of detection (Cryo-EM structure or Nanopore DRS), their helix and domain position on the rRNA, as well as their functional role are presented for each modified residue.

**<u>Table 5 :</u>** List of oligonucleotides

**<u>Table 6 :</u>** Read statistics from Deeplexicon demultiplexing, length filtering and read selection

## Supplementary Movie Legends

**<u>Movie 1 :</u>** Result of the cryoSPARC 3D Flexible refinement

The 4 main different modes of motion computed by the cryoSPARC 3D Flexible refinement are shown.

## Material and Methods

### -Mitochondria and ribosome purification

Mitochondria from cauliflower were purified as previously described (Waltz 2019; Waltz 2020; Waltz 2021). For the chloramphenicol stalled mitoribosomes, mitochondria purification was carried out in the presence of 500 µg/mL chloramphenicol. Mitoribosome purification was conducted as previously^68,69^. In brief, purified mitochondria were re-suspended in Lysis buffer (20 mM HEPES-KOH, pH 7.6, 70 mM KCl, 20 mM MgCl_2_, 1 mM DTT, 1.6% Triton X-100, supplemented with proteases inhibitors (C0mplete EDTA-free)) to a 1.3 mg/ml concentration and incubated for 15 min in 4°C. Lysate was clarified by centrifugation at 25,000 g, 20 min at 4°C. The supernatant was loaded on a 40% sucrose cushion in Monosome buffer (Lysis buffer with 0.1% Triton X-100) and centrifuged at 235,000 g, 3h, 4°C. The crude ribosome pellet was re-suspended in Monosome buffer and loaded on a 10-30 % sucrose gradient in the same buffer and ran for 16 h at 65,000 g. Fractions corresponding to mitoribosomes were collected, pelleted and re-suspended in Monosome buffer and directly plunge frozen for cryo-EM analysis. For the samples treated with chloramphenicol or GTP analog the same procedure was used, but with 200µg/mL chloramphenicol or 200 μM Guanosine-5’-(γ-thio)-triphosphate, Tetralithium salt, (GTPγS).

### -Preparation of total RNA and DNA from purified mitochondria

Total mitochondrial RNA was extracted from purified cauliflower mitochondria pellet using TRI reagent (Sigma-Aldrich). After isopropanol precipitation, the RNA pellet was washed with 70% ethanol, air-dried, and resuspended with nuclease-free water. Total RNA was then treated with Turbo DNase (Thermo Fisher Scientific), purified with phenol/chloroform, and further subjected to RNA Clean and Concentrator kit (Zymo Research). The total mitochondrial RNA was used for direct RNA sequencing (DRS) library preparation. Total mitochondrial DNA was extracted from purified cauliflower mitochondria pellet using TRI reagent (Sigma-Aldrich). The sequences of 26S and 18S rRNA genes were introduced in a plasmid vector, that was used as a template for their subsequent amplification. The rRNA genes were amplified with primers 26S_F/26S_R and 18S_F/18S_R for 26S and 18S respectively. The sequence of the T7 polymerase was introduced to the 5’ of the amplified products via their forward primer. *In vitro* transcription was performed using the T7 RiboMAX Large Scale System (Promega), according to the manufacturer’s instructions. The reactions were subsequently treated with DNase (Promega), purified with phenol/chloroform, and further subjected to MicroSpin G-50 columns (Cytiva). The *in vitro* transcribed 26S and 18S rRNAs were used for DRS library preparation.

### -Direct RNA sequencing library preparation

For DRS library preparation, custom reverse transcription adapters (RTAs) containing Deeplexicon multiplexing barcodes (BC1, BC2 or BC3)^70^ were designed for sequence-specific ligation to the 3’-ends of *B. oleracea* mitochondrial rRNAs (Supplementary Table 5). For native rRNAs, BC1 was included in the RTA oligos (Oligo A and Oligo B), whereas for each IVT rRNA a different barcode sequence was assigned: BC3 for IVT 26S and BC2 for IVT 18S. Oligo A and Oligo B for each rRNA target were annealed (1:1) at 2 mM in annealing buffer (10 mM Tris-HCl pH 7.5, 50 mM NaCl) by heating to 95°C for 2 min and letting them cool down gradually (0.1°C/sec). The libraries were prepared following the Oxford Nanopore Technologies (ONT) DRS protocol for PromethION FLO-PRO002 R9.4.1 flowcells using reagents from the DRS kit (SQK-RNA002, ONT) and the custom RTAs. Briefly, 1 μg of total mitochondrial RNA or 2 pmol of each IVT rRNA were ligated with their respective custom RTA using T4 DNA ligase (Thermo Fisher Scientific) for 15 min at room temperature, before they were reversed transcribed with SuperScript III Reverse Transcriptase (Thermo Fisher Scientific). The RNA:DNA hybrid products were purified with 1.8× Agencourt RNAClean XP beads (Thermo Fisher Scientific), washed with 70% ethanol, and ligated with the RNA Adapter (RMX). The products were purified with 0.4× Agencourt RNAClean XP beads, washed twice with wash buffer (WSB), and eluted in elution buffer. Finally, the RNA library was mixed with RNA Running Buffer (RRB) and loaded onto a primed FLO-PRO002 flowcell (ONT).

### -Basecalling and demultiplexing of ONT direct RNA sequencing data

The ONT output from three DRS runs performed on two distinct FLO-PRO002 flowcells (Supplementary Table 6) were processed via *deeplexicon_sub*.*py* dmux (Deeplexicon v1.2.0, options: - f multi -s 0.95 -m models/resnet20-final.h50)^70^ on an Nvidia V100 GPU accelerator to obtain demultiplexing calls for all the fast5 reads. These reads were basecalled with Guppy (v6.5.7, config: “rna_r9.4.1_70bps_hac_prom.cfg”) and the fastq output split into separate files (BC1, BC2, BC3, BC4). The demultiplexed and basecalled ONT reads were filtered by length (2000-3300 nt for 26S and 1700- 2000 nt for 18S reads) and then aligned to the *B. oleracea* rRNA reference sequences in Geneious (v2024.0.7) using “Map to Reference” with the built-in mapper. Near full-length reads were selected for current intensity analysis to determine whether nucleotide modifications observed by cryo-EM could be validated by ONT DRS.

### -Analysis and visualization of current intensities

Raw fast5 files were basecalled again using Guppy (v6.3.8, options: “--kit SQK-RNA002 --flowcell FLO-PRO002 –fast5_out”) in order to include FastQ entries, Move and Trace tables, which are necessary for the current intensity analysis. Full length reads were selected from the fastq file and mapped with minimap2 (v2.26, options: “-ax splice -uf -k14”) to their corresponding reference sequences. <u>T</u>o extract event-level information from these basecalled fast5 files, we used f5c (v1.5, https://github.com/hasindu2008/f5c)^71^, a GPU-accelerated implementation of Nanopolish (https://github.com/jts/nanopolish) that aligns ONT signals called “events” to reference k-mers. First, we indexed the FastQ file using the command *f5c index* to link each read to its basecalled fast5 file. Next, we extracted per-read current intensities using *f5c eventalign* with the options “--rna --print-read-names --collapse-events -b $bam -g $ref_sequence -r $fastq > f5c_eventalign.tsv”. The resulting alignment and eventalign output files were then analyzed in R. Namely, we used the *pileup* function from the Rsamtools package (v2.20.0) to obtain nucleotide frequencies for each position in the alignments. We then incorporated eventalign information to display the nanopore-derived current intensities at specific nucleotide positions. All visualizations were performed using ggplot2 (v3.5.1).

### -Cryo-EM grid preparation

4 µL of purified ribosome at a protein concentration (OD280) of 3 µg/µL was applied onto Quantifoil R2/1 200-mesh holey carbon grids, pre-coated with a 2 nm continuous carbon film and glow-discharged (2.5 mA for 20 sec). Samples were incubated on the grid for 25 sec and then blotted with filter paper for 2 sec in a temperature and humidity controlled Vitrobot Mark IV (T = 4°C, humidity 100%, blot force 5) followed by vitrification in liquid ethane.

### -Single particle cryo-EM Data collection

Data collection was performed using a 300kV G4 Titan Krios electron microscope (ThermoScientific) equipped with Falcon4i camera and a Selectris X energy filter using EPU for automated data acquisition at the IGBMC’s Integrated Structural Biology platform. Data were collected at a nominal underfocus of –0.5 to −2.5 µm at a magnification of 165,000x, yielding a pixel size of 0.729 Å.

For the monosomes, two datasets (1 and 2) were recorded as movie stacks, from 2 different grids of 9,091 and 7,825 micrographs respectively and exported as EER files. Each movie stacks were fractionated into 40 frames for a total electron dose of 41.66 e^−^/Å^2^. For the small subunit dataset, 20,145 micrographs were recorded as movie stacks and exported as EER files. Each movie stacks were fractionated into 40 frames for a total electron dose of 50.96 e^−^/Å^2^.

For the stalled monosomes with P-site tRNA, data collection was performed using a 300kV Titan Krios electron microscope (ThermoScientific) equipped with a K2 camera and a GIF energy filter using SerialEM for automated data acquisition at the Biozentrum’s BioEM facility. Data were collected at a nominal underfocus of –0.5 to −2.5 µm at a magnification of 130,000×, yielding a pixel size of 1.058 Å. 5,935 micrographs were recorded as movie stacks, exported as TIF files. Each movie stacks were fractionated into 50 frames for a total electron dose of 46 e^−^/Å^2^.

### -Single particle cryo-EM Data processing

For all datasets, the entire processing pipeline was carried out in cryoSPARC^72^ and the processing workflows are shown Supplementary Fig. 1, 2, 3 and 4.

For the monosome datasets, pre-processing and particle picking were performed independently and the resulting good particles were then merged. For dataset 1, after motion correction and CTF estimation 8,829 micrographs were kept, and 7,532 micrographs were kept for dataset 2. For particle picking and 2D classification, the same approach used for the small subunit dataset was used but with particles extracted with a box size of 756 pixels, down sampled to 256 pixels (pixel size of 2.1528). For dataset 1, 137,759 particles remained after 2D classification and then 111,531 after initial 3D classification. For dataset 2, 205,994 particles remained after 2D classification and then 174,386 after initial 3D classification. At this point, particles were merged, resulting in a total number of 258,917 particles that were further cleaned down to 219,550 particles. They were re-extracted for high resolution refinement, using a box size of 756 pixels, down sampled to 560 pixels (pixel size of 0.9841). A global refinement reached 2.77 Å resolution. After Global and Local CTF refinement, resolution improved to 2.34 Å. Reference Based Motion Correction was then performed and further improved the resolution to 2.18 Å. Another round of Global and Local CTF refinement at full resolution (bin1) improved the resolution to 2.11 Å. Particles were down sampled to 588 pixels (pixel size of 0.9373), and focused refinement were performed using a mask for the LSU (1.99 Å), and then further local refined for the LSU L7/12 Stalk (2.85 Å), the Central Protuberance (2.09 Å) and the LSU’s back (2.53 Å). The same was done for the SSU (2.35 Å), and then SSU head (2.24 Å), head protuberance (2.41 Å), SSU body (2.25 Å), foot (2.47 Å) and body protuberance (2.58 Å). To improve the SSU head extension, signal subtraction (with a mask to remove the signal of most of the ribosome) and particle recentering was performed, allowing to resolve the head extension to 3.87 Å and then further to 3.43 Å for the base, 4.04 Å for the core and 4.32 Å for the tip.

For the small subunit dataset, movies were motion corrected and CTF was estimated using Patch Motion Correction and Patch CTF. Bad micrographs were discarded, resulting in 19,932 out of 20,145 micrographs remaining. Particles were then picked using a template picker approach, yielding 1,896,051 initial positions that were extracted with a box size of 720 pixels, down sampled to 280 pixels (pixel size of 1.8746) for faster processing. Particles were cleaned iteratively through 2 rounds of 2D classification from which 931,475 particles were selected. Bad particles were then further discarded in 3D through a combined approach of Ab-initio, Heterogeneous Refinement and 3D Variability analysis, from which 602,798 particles remained. Good particles were then further sorted using a mask covering the body of the SSU, resulting in 2 main classes: an intact empty small subunit class (388,094 particles), and the RsgA class (208,855 particles). At this point, particles were re-extracted for high resolution refinement, using a box size of 720 pixels, down sampled to 560 pixels (pixel size of 0.9373). A global refinement of the empty small subunit class reached 2.3 Å resolution while the RsgA class reached 2.44 Å resolution. After Global and Local CTF refinement, resolution improved to 2.05 Å and 2.25 Å respectively. Reference Based Motion Correction was then performed and further improved the resolution to 2.04 Å and 2.27 Å respectively. Another round of Global and Local CTF refinement was performed at bin1. Particles were down sampled to 588 pixels (pixel size of 0.8927), and then focused refinements were performed using masks for the SSU head and body and for RsgA resulting in, for the RsgA class, 2.07 Å for the RsgA focused map, 2.17 Å for the body and 2.07 Å head.

For the stalled monosomes, 5,621 micrographs were kept after motion correction and CTF estimation. Particles were picked and extracted with a 564 pixels’ box size, down sampled to 282 pixels (pixel size of 2.1160) and cleaned through 2 rounds of 2D classification resulting in 295,205 particles. After cleaning of the particles in 3D, 219,458 particles were selected and submitted to focused 3D classification with a mask covering the decoding center, made from A, P and E site tRNA models. This yielded a clear class of P-site tRNA stalled monosomes with 34,964 particles. For high resolution refinement, particles were re-extracted using a box size of 500 pixels (pixel size of 1.1934). After Global and Local CTF refinement the resolution reached 3.17 Å. Reference Based Motion Correction was then performed and further improved the resolution to 3.05 Å. Another round of Global and Local CTF refinement at full resolution (bin1) and then focused refinements were performed using masks for the LSU and SSU that respectively reached 2.95 Å and 3.08 Å.

### -Model building and refinement

Even though a model for the plant mitoribosome was already available (PDB: 6XYW), we decided to build all the proteins *de novo*, taking a completely unbiased approach and making the most of these high-resolution maps. For that, focused refined cryo-em maps (all at resolution better than 3 Å) for the different main parts of the mitoribosome (large subunit, body and head of the small subunit) were used as inputs for ModelAngelo^32^ without input sequence. Peptide chains obtained were then blasted against the Brassicaceae taxon (3700) of the UNIPROT database, resulting mainly on hits from *Arabidopsis thaliana*, and *Brassica oleracea var. oleracea* and the corresponding AlphaFold2 models were retrieved for all the identified proteins and then matched to the ModelAngelo models using the Matchmaker tool in ChimeraX^73^, which were further rigid-body fitted in their respective cryo-EM maps. The protein models were then manually inspected and refined in COOT^74^, where the best fitting protein, sequence-wise, were chosen. For the rRNAs, the models from 6XYW were rigid-body fitted in the density and then refined with restraints in COOT. rRNA extensions in lower resolutions areas (e.g., the head extension of the SSU or back extension of the LSU) were modeled just as stretches of A and U. The sequences were then mutated accordingly and refined again. Ions, ligands and rRNA modifications were first placed by homology with *E. coli* 8B0X and then manually curated in COOT. Finally, water molecules were placed using the PHENIX phenix.douse tool and curated using the “Check Waters” tool in COOT. The different parts of the model were then automatically refined in PHENIX^75^ against the best resolved focus-refined maps using the phenix.real_space_refine tool and then again manually refined in COOT, this repeating through several cycles. Chimeric maps were generated by using the Map Box option in PHENIX to cut out densities around the models (3.5 Å around the atoms). Maps were then fitted into the consensus maps in ChimeraX and combined using the ‘vop maximum’ command. The model geometry was validated using MolProbity^76^.

## -Figure preparation

All figures were prepared using ChimeraX^77,78^.

## Author contributions

VS, TTN and NC performed biochemical purifications. FW and VS performed cryo-EM data processing, model building and interpretation, and initial writing of the manuscript. VS, DP, and TB performed nanopore analyses. FW and PG conceived the project, interpreted results, and finalized the manuscript.

## Acknowledgments

We thank M. Chami and D. Kalbermatter from the University of Basel - Biozentrum BioEM facility for their help in operating the electron microscopes. We thank Prof. Dr. Benjamin D. Engel for hosting Dr. Florent Waltz at the Biozentrum. We thank Dr. Asier González Seviné and Prof. Dr. Mihaela Zavolan for their access to the BioComp gradient fractionator and Dr. Irene Vercellino for advice on generating chimeric cryo-EM maps. This work used the Integrated Structural Biology platform of the Strasbourg Instruct-ERIC center IGBMC-CBI supported by FRISBI (ANR-10-INBS-0005). We thank Dr. Alexandre Durand for his help in operating the electron microscopes. We thank the Strasbourg Esplanade proteomic platform for the proteomic analysis.

This work was supported by the “Centre National de la Recherche Scientifique”, the University of Strasbourg, by Agence Nationale de la Recherche (ANR) grants ‘DAMIA, ANR-20-CE11-0021’ and ‘PROPHAN, ANR-22-CE12-0008-01’ to PG and by the LabEx consortium “MitoCross” in the frame of the French National Program “Investissement d’Avenir” ANR-11-LABX-0057_MITOCROSS. This work was supported by the Swiss National Science Foundation (SNSF) with a Swiss Postdoctoral Fellowships (Grant number 210561) and by the Alexander von Humboldt Foundation through the Humboldt Research Fellowship Programme for Postdocs to FW.

## Data Availability

The single particle cryo-EM maps of *B. oleracea* mitoribosome have been deposited at the Electron Microscopy Data Bank (EMDB) and models on the protein data bank (PDB). Full high-resolution mitoribosome EMD-51718 (PDB 9GYT), unfocused EMD-51703, focused LSU EMD-51710, focused LSU CP EMD-51711, focused LSU L7/12 stalk EMD-51712, focused LSU back extension EMD-51713, focused LSU rPP5 EMD-51714, focused SSU body EMD-51704, focused SSU body protuberance EMD-51709, focused SSU foot rPPR1 EMD-51708, focused SSU foot rPPR10 EMD-51707, focused SSU head EMD-51705, focused SSU head S3 area EMD-51706, focused SSU head extension base EMD-51715, focused SSU head extension core EMD-51716, focused SSU head extension tip EMD-51717. Small subunit in presence of RsgA EMD-50014 (PDB 9EVT), focused SSU head EMD-50015, focused SSU body EMD-50017, focused RsgA EMD-50016. P-site tRNA mitoribosome stalled with chloramphenicol EMD-50011 (PDB 9EVS), focused LSU EMD-50012, focused SSU EMD-50013.

