## Supplementary Fig. 2 for "Structural insights into maturation and translation of a plant mitoribosome"

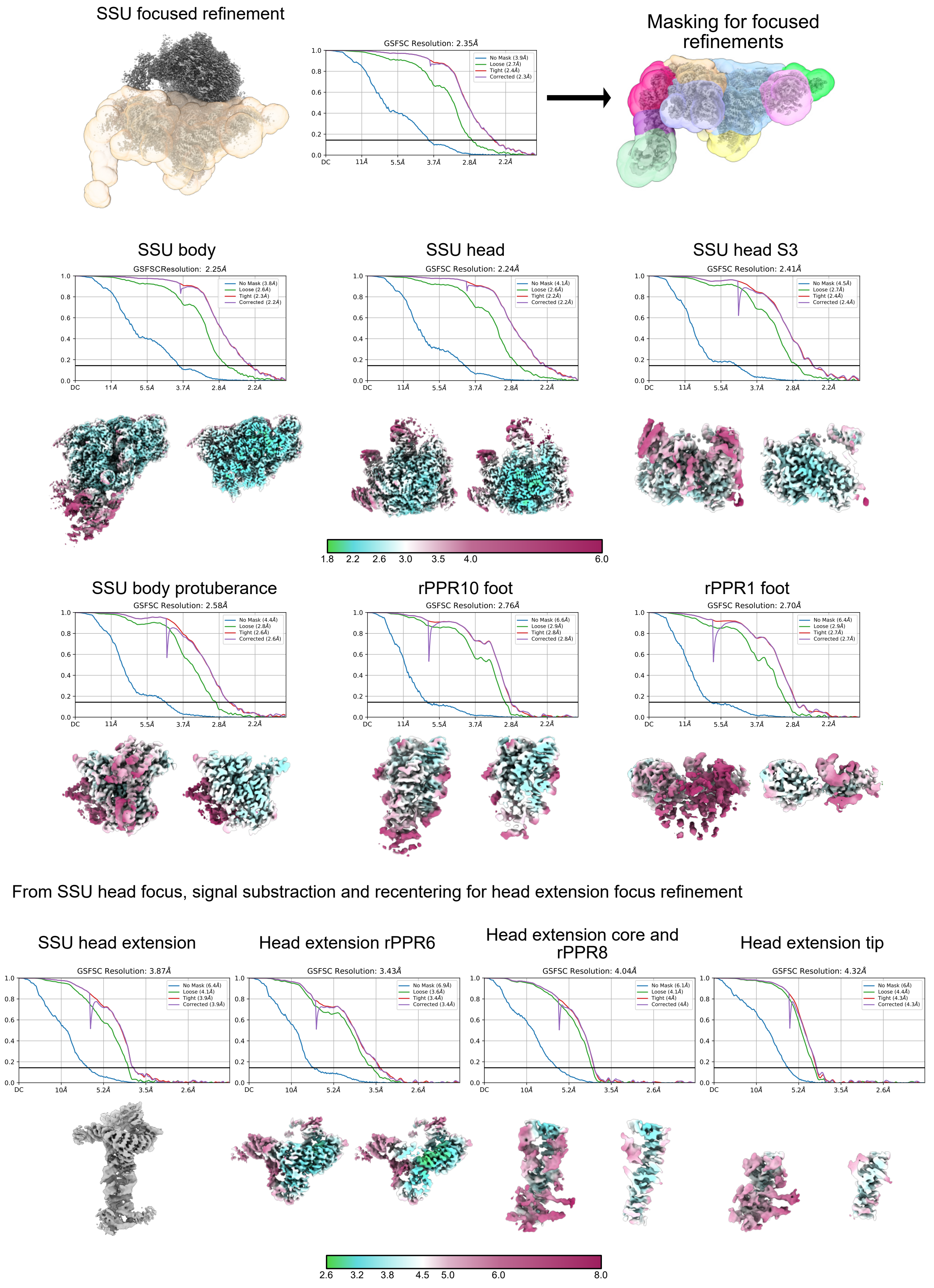


**Supplementary Figure 2 :** FSC plots of the SSU from the high resolution mitoribosome

Focused refinements for the SSU. For the final reconstruction, GSFSC curves are plotted, with resolution calculated at the 0.143 threshold. Local resolution plotted on the maps were generated using the built-in cryoSPARC tool with default parameters (FSC=0.5 threshold), with a resolution scale different for the SSU head extension, with maps also shown in cut view.
