## Supplementary Fig. 3 for "Structural insights into maturation and translation of a plant mitoribosome"

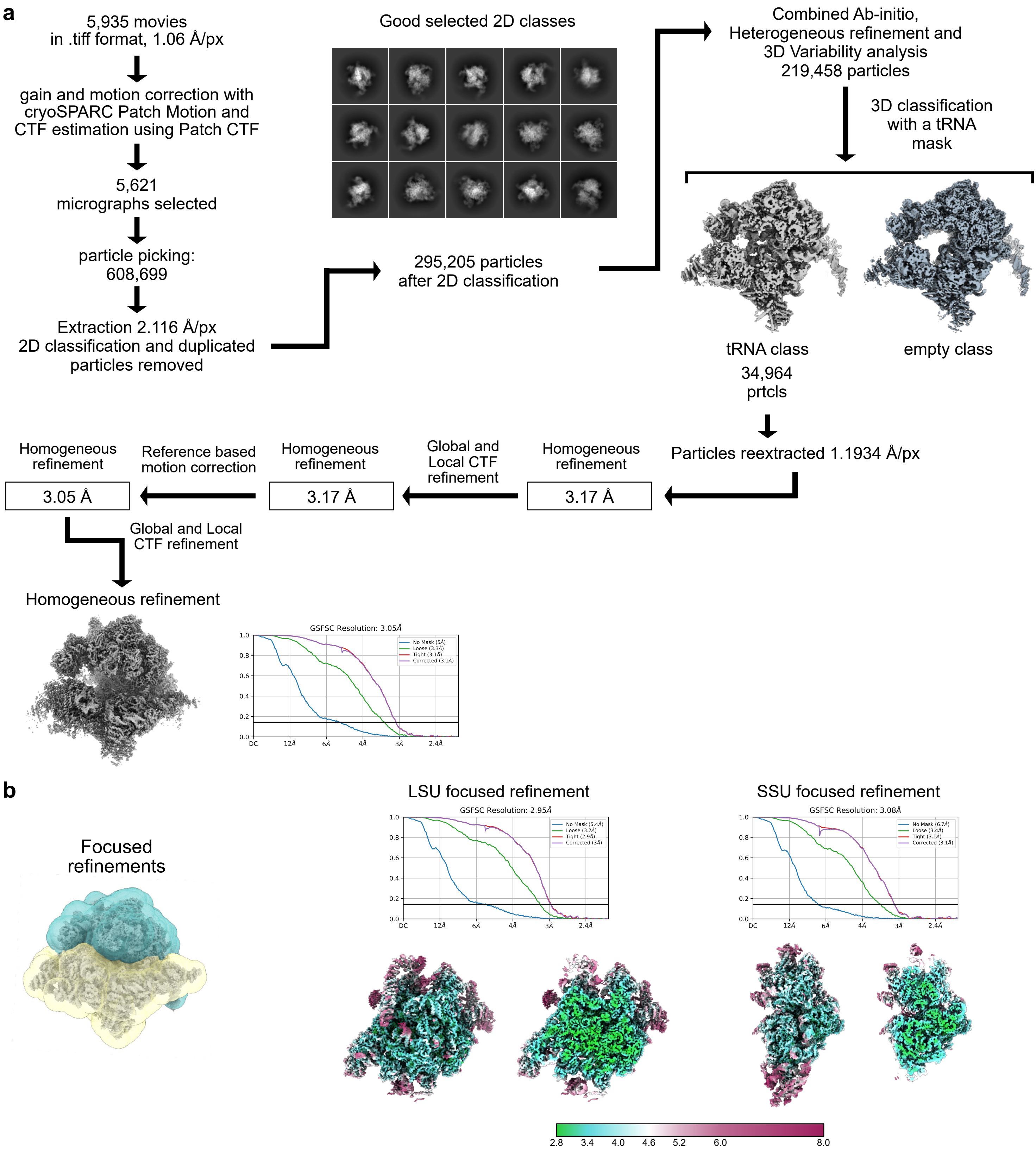


**<u>Supplementary Figure 3 :</u>** Single-particle data processing workflow of the stalled tRNA mitoribosome

Graphical summary of the processing workflow described in Methods. **a** Post-processing, 2D and 3D classification. **b** Focused refinements for the tRNA class. For the final reconstruction, GSFSC curves are plotted, with resolution calculated at the 0.143 threshold. Local resolution plotted on the maps were generated using the built-in cryoSPARC tool with default parameters (FSC=0.5 threshold), all using the same resolution scale, with maps also shown in cut view.
