## Supplementary Fig. 5 for "Structural insights into maturation and translation of a plant mitoribosome"

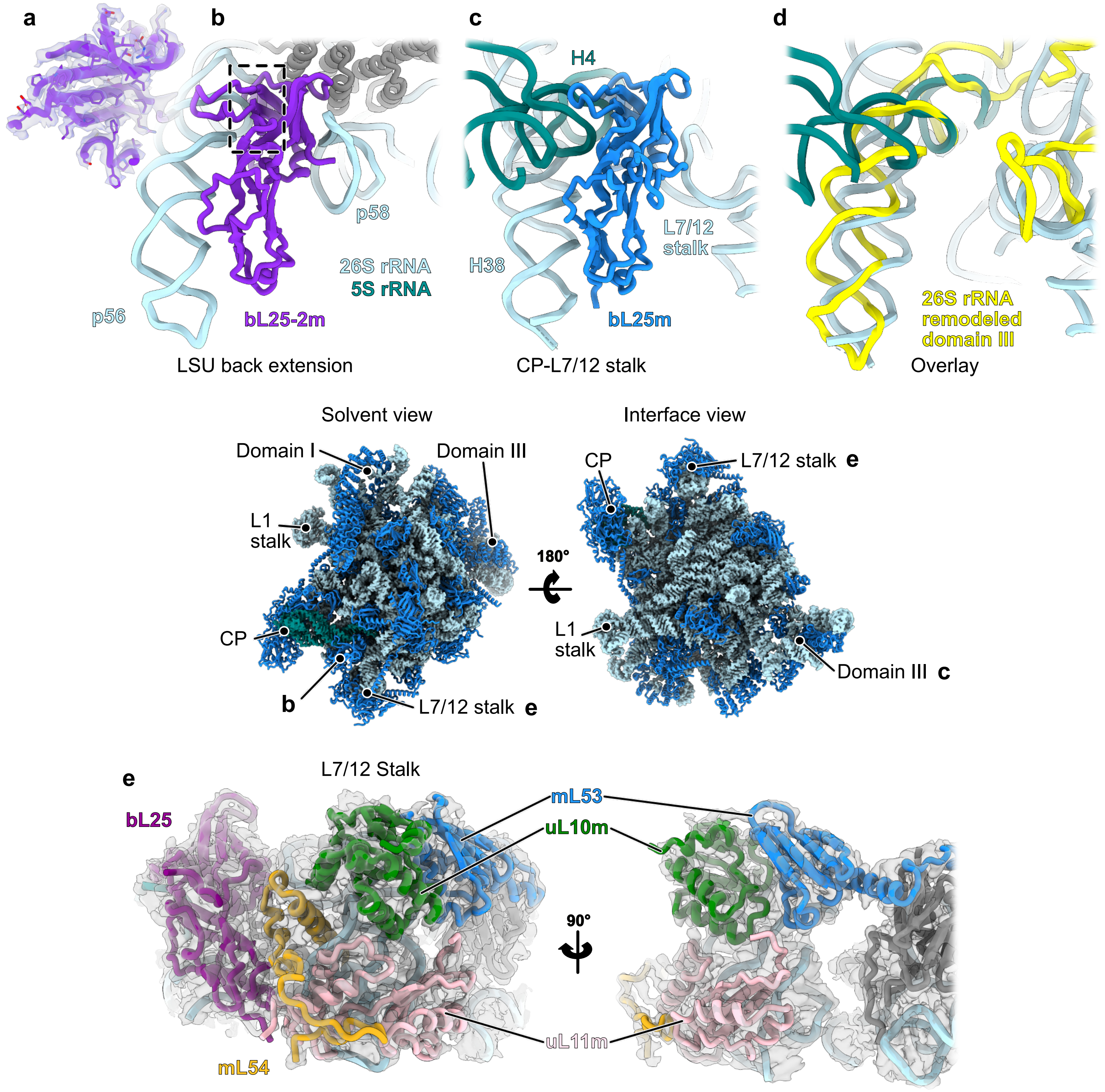


**Supplementary Figure 5 :** Newly resolved r-proteins in the LSU

**a-d** Newly identified r-protein bL25-2m in the LSU back extension, remodeling the 26S domain III. **a** show a close-up of bL25-2m, shown in **b**, with the experimental density. **c** Close-up view of the canonical bL25m r-protein in the mitoribosome. **d** Overlay of the remodeled domain III from **b**, shown in yellow, with the 5S and 26S rRNA from **c** showing the structural mimicry of the rRNA, with the base of p56 mimicking H4 of the 5S and p56's tip mimicking H38, as well as p58 mimicking the base of the L7/17 stalk. **e** View of the L7/12 stalk with the experimental density showing mL54 which was previously unresolved. The different areas of the LSU are shown in the middle of the figure, with panel **b**, **c** and **e** positions indicated.
