## Supplementary Fig. 6 for "Structural insights into maturation and translation of a plant mitoribosome"

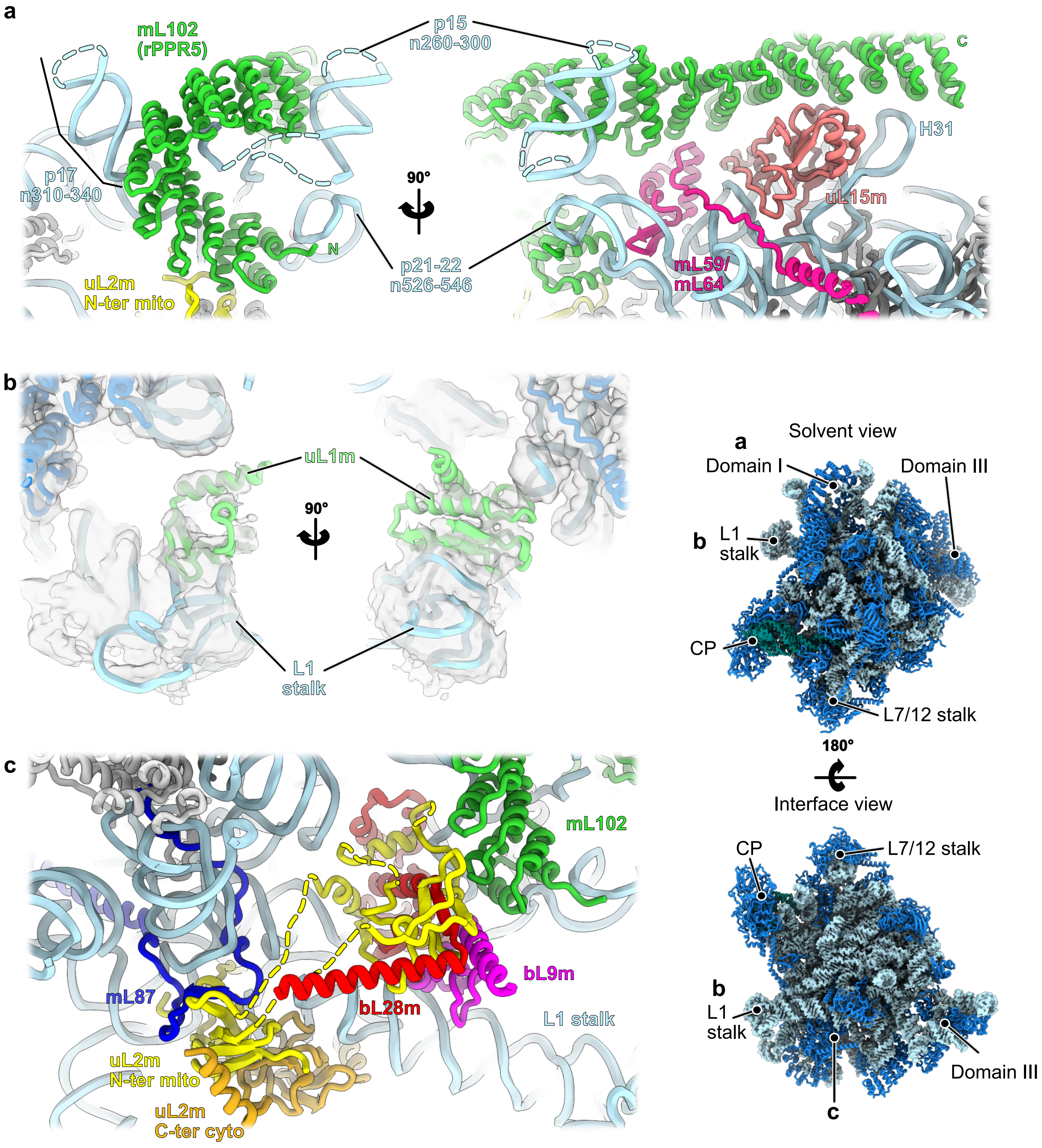


**<u>Supplementary Figure 6 :</u>** Improved model in the LSU

**a** Close-up view of the mL102 area. The rPPR protein mL102 interacts with three r-proteins, uL15m, mL59/64 and uL2m (N-ter mito). The N-terminal part of the protein largely interacts with rRNA and contributes to the remodeling of domain I, forming p17, p17 and p21-22, specific helices of the plant mitoribosome. Unmodeled densities of the rRNA are shown are blue dashed lines. **b** View of the L1 stalk at low threshold, revealing the presence of the uL1m protein. **c** In the plant mitoribosome, uL2m is split into two parts. What would correspond to the N-terminal part of the canonical uL2 is encoded as uL2m in the mitochondrial genome (yellow), whereas the C-terminal part is encoded in the nuclear genome (orange). The uL2m N-ter mito possesses large insertions that extend away from the uL2 core to make intricate interactions with bL28m, bL9m and mL102. Close to the uL2 core it also makes direct interactions with mL87. The different areas of the LSU are shown on the right hand-side of the figure, with panel **a**, **b** and **c** positions indicated.
