## Supplementary Fig. 7 for "Structural insights into maturation and translation of a plant mitoribosome"

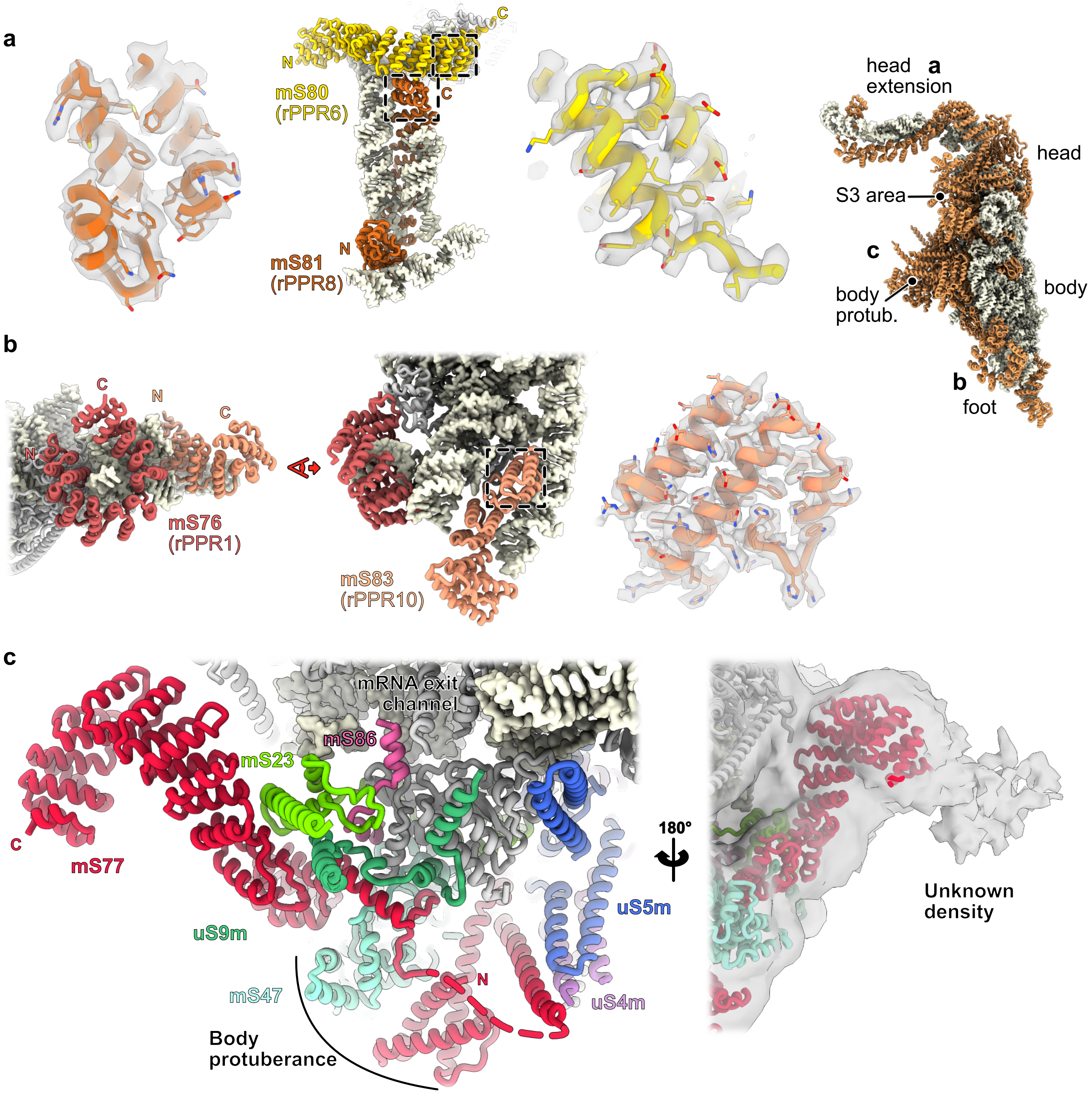


**Supplementary Figure 7 :** PPR proteins of the SSU

**a** On the SSU head extension, two PPR proteins are found, mS80 and mS81. Portions of the atomic models (highlighted by dashed boxes) are shown in the experimental density. **b** At the foot of the SSU, two PPR proteins are found, m76 and mS83. Portions of the atomic models (highlighted by dashed boxes) are shown in the experimental density. **c** The newly identified r-protein mS77 (red) spans from the SSU body protuberance to the exit mRNA channel. It is composed of two main domains. Forming part of the SSU body protuberance, the N-terminal part interacts with extensions of uS5m and uS4m as well as mS47. Through a flexible portion of the protein (represented by dashed-lines) it wraps around the SSU body and its C-terminal part is composed of a long stretch of PPR repeats, interacting with mS23 and an extension of uS9m. At the mRNA exit channel a small portion of mS86 is also visible, sandwiched between mS23 and uS2m. At lower threshold, we also observe additional densities at the C-terminus of mS77, that could indicate that it serves as a binding platform for additional factors. C- and N-termini of the proteins of interest are indicated by colored N and C. The different areas of the SSU are shown on the right hand-side of the figure, with panel **a**, **b** and **c** positions indicated.
