## Supplementary Fig. 8 for "Structural insights into maturation and translation of a plant mitoribosome"

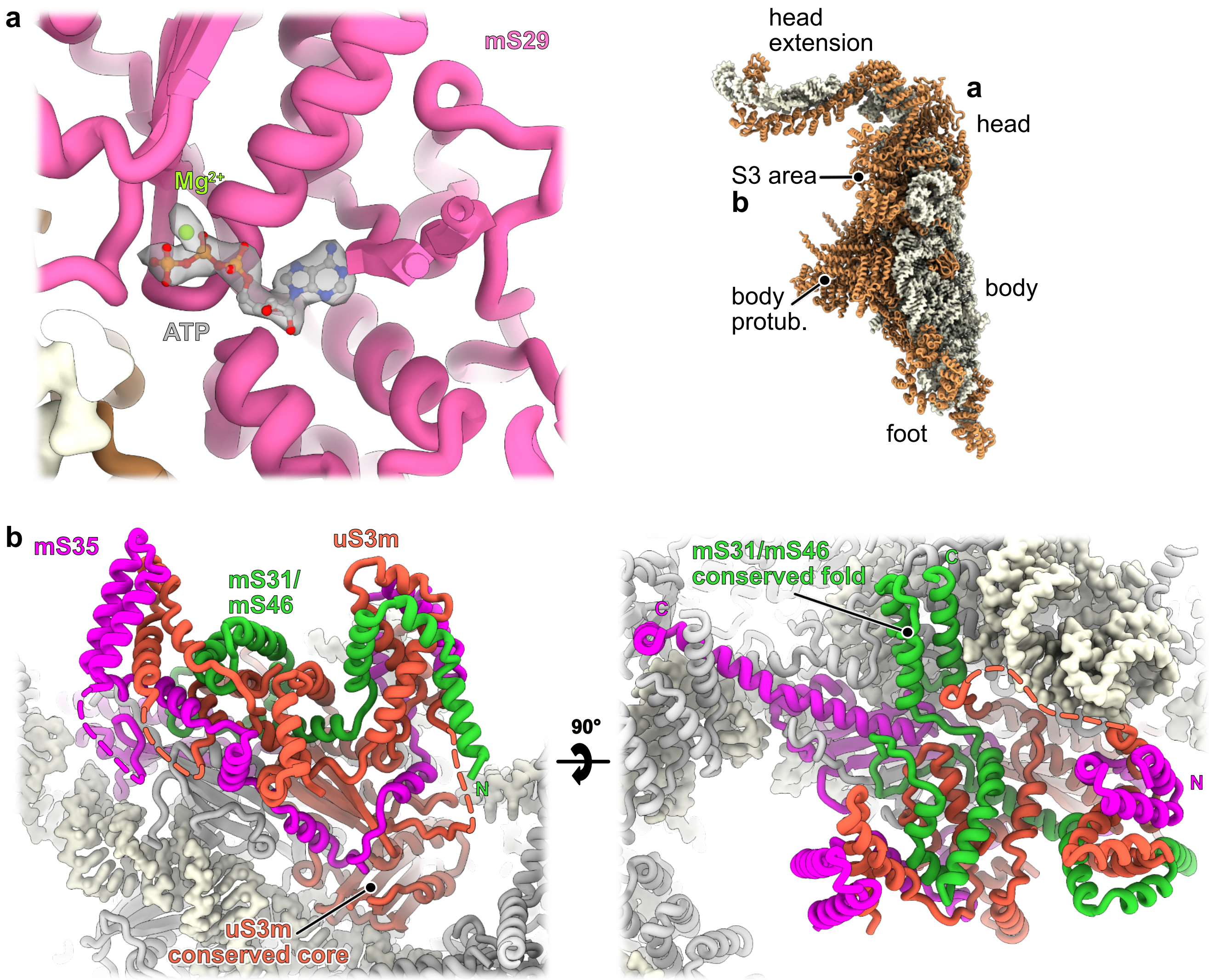


**Supplementary Figure 8 :** Improved model in the SSU

**a** Close-up view of mS29 on the SSU head, showing the ATP with $Mg^{2+}$ in its density. **b** View of the SSU head protuberance, or S3 area. This domain is formed by a large insertion in uS3m, forming intricate interactions with the N-terminal part of mS35 and a large portion of mS31/mS46. with C- and N-termini of the proteins of interest are indicated by colored N and C. The different areas of the SSU are shown on the right hand-side of the figure, with panel **a** and **b** positions indicated.
