## Supplementary Fig. 9 for "Structural insights into maturation and translation of a plant mitoribosome"

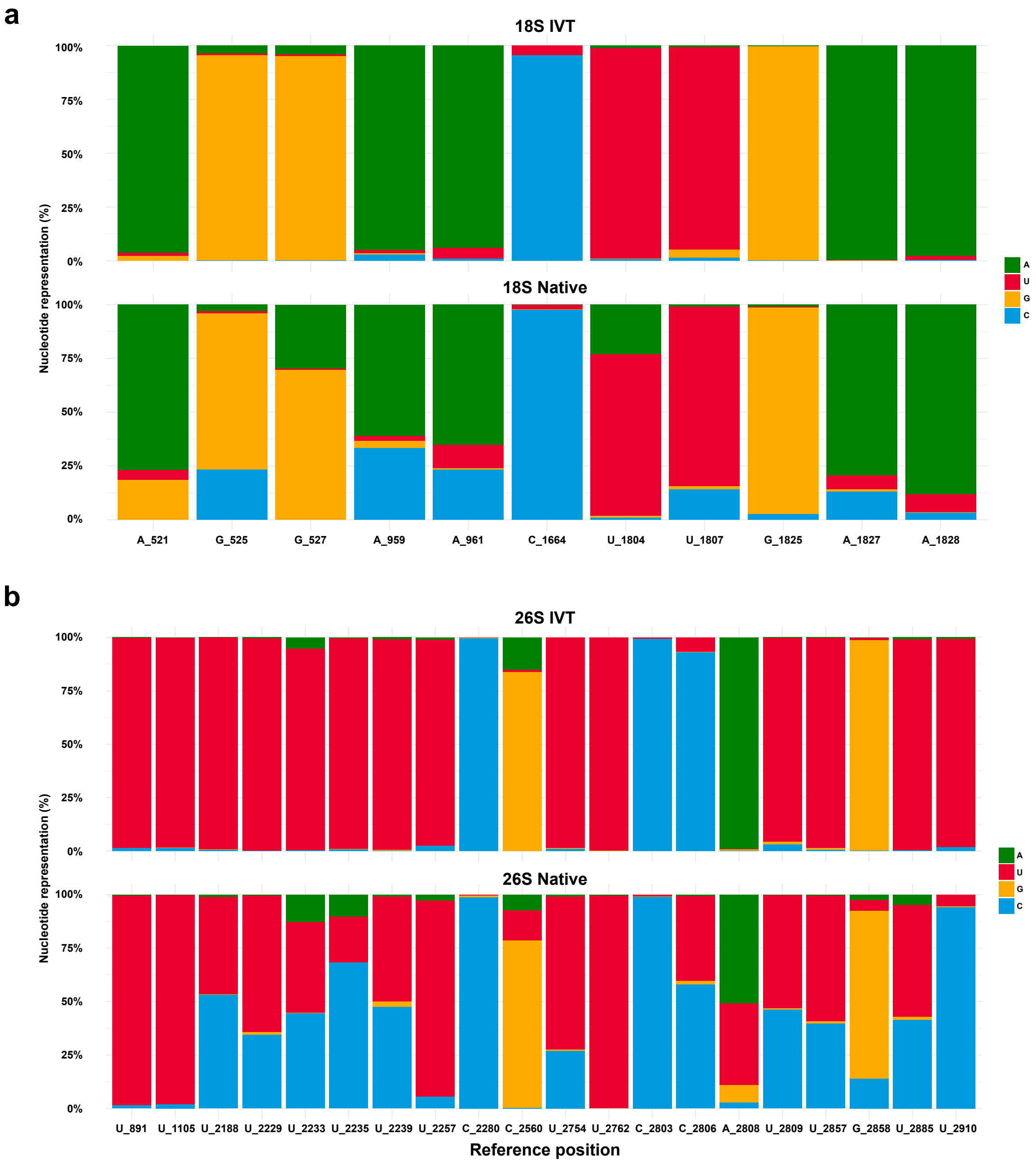


**Supplementary Figure 9 :** Nucleotide frequency at positions with detected modifications

Base-called nucleotides and their percentage presented for all the positions with detected modifications of 18S rRNA (**a**) and 26S rRNA (**b**). Comparison of IVT and Native rRNAs is presented, with the latter displaying base-calling errors accompanying the presence of rRNA modifications. Positions with identified modifications from the cryo-EM map are also included. Base-called nucleotides are colored.
