## Supplementary Fig. 10 for "Structural insights into maturation and translation of a plant mitoribosome"

**a**

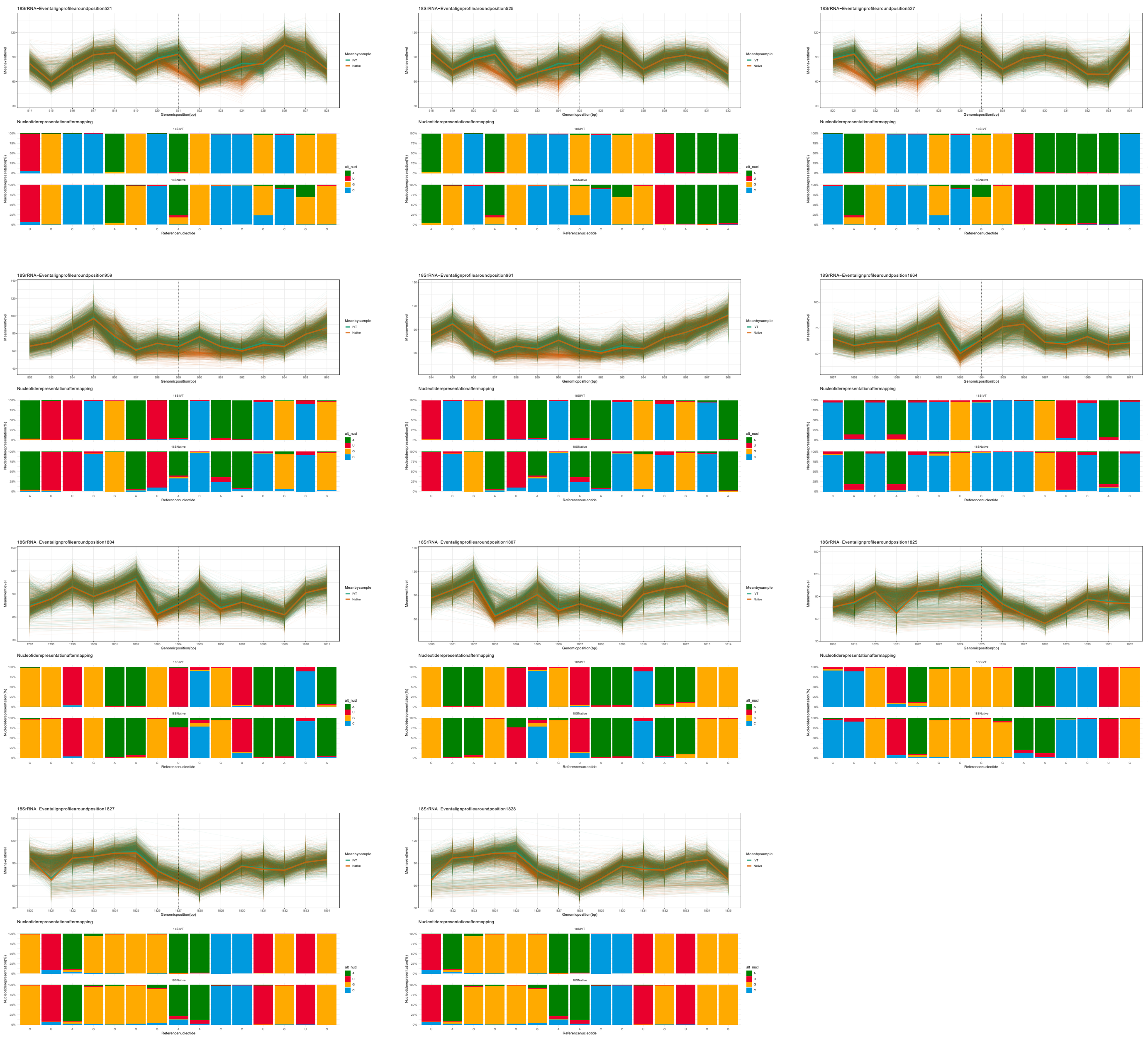
18SrRNA−Eventalignprofilearoundposition521
18SrRNA−Eventalignprofilearoundposition525
18SrRNA−Eventalignprofilearoundposition527
18SrRNA−Eventalignprofilearoundposition959
18SrRNA−Eventalignprofilearoundposition961
18SrRNA−Eventalignprofilearoundposition1664
18SrRNA−Eventalignprofilearoundposition1804
18SrRNA−Eventalignprofilearoundposition1807
18SrRNA−Eventalignprofilearoundposition1825
18SrRNA−Eventalignprofilearoundposition1827
18SrRNA−Eventalignprofilearoundposition1828
Meanbysample
IVT
Native
Genomicposition(bp)
Nucleotiderepresentationaftermapping
alt_nucl
A
U
G
C
Referencenucleotide

**b**

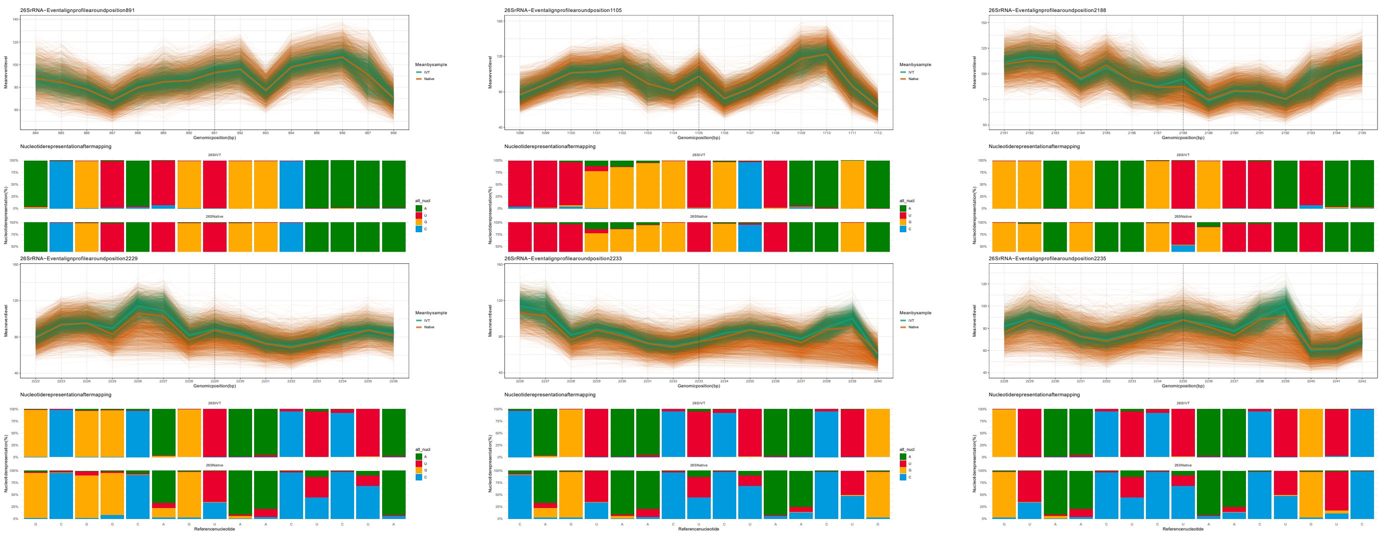
26SrRNA−Eventalignprofilearoundposition891
26SrRNA−Eventalignprofilearoundposition1105
26SrRNA−Eventalignprofilearoundposition2188
26SrRNA−Eventalignprofilearoundposition2229
26SrRNA−Eventalignprofilearoundposition2233
26SrRNA−Eventalignprofilearoundposition2235
Meanbysample
IVT
Native
Genomicposition(bp)
Nucleotiderepresentationaftermapping
alt_nucl
A
U
G
C
Referencenucleotide

**b**

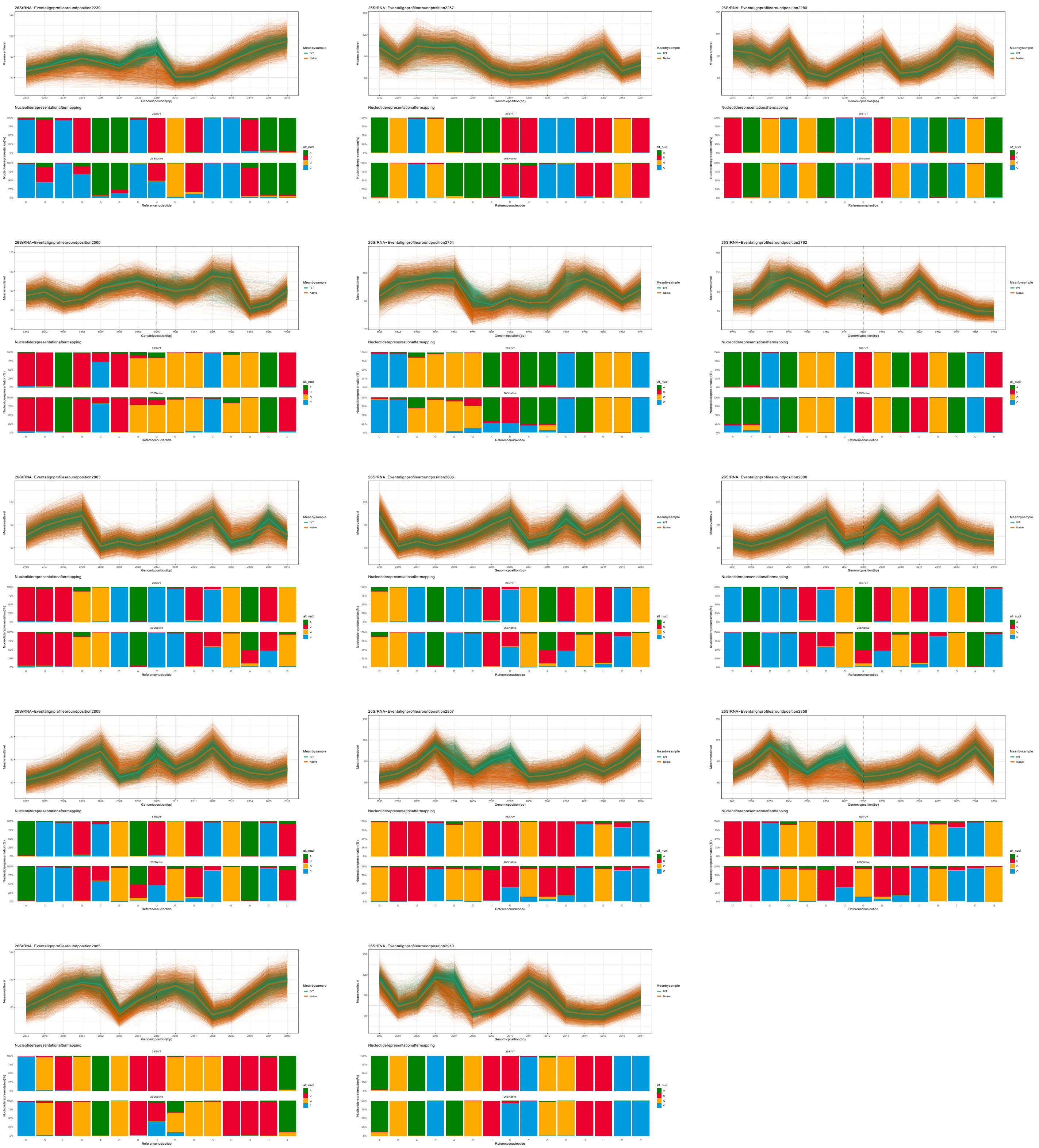


**Supplementary Figure 10 :** Current intensity and nucleotide frequency at modified positions

Per-read current intensity analysis and nucleotide frequency in the 15-mer regions surrounding the modified sites of 18S rRNA (**a**) and 26S rRNA (**b**). Green lines correspond to native reads whereas brown lines to IVT reads. Thick green and brown lines represent the mean current intensity of native and IVT reads, respectively.
