## Supplementary Table 1 for "Structural insights into maturation and translation of a plant mitoribosome"

| EMDB | Unfocused | LSU | LSU CP | LSU L7/12 | LSU extension | LSU rPPR5 | SSU body | SSU body protub. | SSU foot rPPR1 | SSU foot rPPR10 | SSU head | SSU head S3 | SSU head ext. base | SSU head ext. core | SSU head ext. tip |
|---|---|---|---|---|---|---|---|---|---|---|---|---|---|---|---|
| | Full high resolution (PDB 9GYT - EMD-51718) | | | | | | | | | | | | | | |
| EMDB | EMD-51703 | EMD-51710 | EMD-51711 | EMD-51712 | EMD-51713 | EMD-51714 | EMD-51704 | EMD-51709 | EMD-51708 | EMD-51707 | EMD-51705 | EMD-51706 | EMD-51715 | EMD-51716 | EMD-51717 |
| **Data collection and processing** | | | | | | | | | | | | | | | |
| Magnification | | | | | | | | 165,000x | | | | | | | |
| Voltage (kV) | | | | | | | | 300 | | | | | | | |
| Electron exposure (e–/Å$^2$) | | | | | | | | 41.66 | | | | | | | |
| Defocus range (μm) | | | | | | | | -0.5 to -2.5 | | | | | | | |
| Pixel size (Å) | | | | | | | | 0.729 | | | | | | | |
| Symmetry imposed | | | | | | | | C1 | | | | | | | |
| Initial particle images (no.) | | | | | | | | 613,488 | | | | | | | |
| Final particle images (no.) | | | | | | | | 219,550 | | | | | | | |
| Map resolution (Å) - FSC 0.143 | 2.11 | 1.99 | 2.09 | 2.85 | 2.53 | 2.09 | 2.25 | 2.58 | 2.70 | 2.76 | 2.24 | 2.41 | 3.43 | 4.04 | 4.32 |
| Map resolution range (Å) | 1.6 - 7.0* | 1.6 - 7.8* | 1.6 - 7.7* | 1.6 - 9.6* | 1.6 - 8.8* | 2.0 - 5.7* | 1.6 - 6.9* | 2.3 - 8.3* | 2.3 - 7.8* | 2.3 - 7.5* | 1.6 - 8.2* | 2.2 - 7.4* | 2.7 - 7.9* | 3.3 - 7.9* | 3.6 - 7.4* |
| **Refinement** | | | | | | | | | | | | | | | |
| Initial model used (PDB code) | | | | | | | | AF2 - 6XYW | | | | | | | |
| CC Model vs. Data (mask) | 0.81 | | | | | | | | | | | | | | |
| Map sharpening *B* factor (Å$^2$) | -26.4 | -25.0 | -32.5 | -62.1 | -46.8 | -31.7 | -36.8 | -54.3 | -62.1 | -62.5 | -37.9 | -51.7 | -67.3 | -139.4 | -227.2 |
| Model composition | | | | | | | | | | | | | | | |
| Non-hydrogen atoms | | | | | | | | 233505 | | | | | | | |
| Residues: Protein - Nucleotide | | | | | | | | 15011 - 4806 | | | | | | | |
| Water | | | | | | | | 10050 | | | | | | | |
| Ligands | | | | | | | | ZN: 2<br>K: 84<br>MG: 349<br>ATP: 1 | | | | | | | |
| *B* factors (Å$^2$) | | | | | | | | | | | | | | | |
| Protein | | | | | | | | 0.00/1098.4/69.02 | | | | | | | |
| Nucleotide | | | | | | | | 0.00/440.22/57.04 | | | | | | | |
| Ligand | | | | | | | | 18.37/90.34/48.08 | | | | | | | |
| R.m.s. deviations | | | | | | | | | | | | | | | |
| Bond lengths (Å) | | | | | | | | 0.006 | | | | | | | |
| Bond angles (°) | | | | | | | | 0.989 | | | | | | | |
| Validation | | | | | | | | | | | | | | | |
| MolProbity score | | | | | | | | 1.41 | | | | | | | |
| Clashscore | | | | | | | | 4.57 | | | | | | | |
| Poor rotamers (%) | | | | | | | | 0.01 | | | | | | | |
| Ramachandran plot | | | | | | | | | | | | | | | |
| Favored (%) | | | | | | | | 96.97 | | | | | | | |
| Allowed (%) | | | | | | | | 2.92 | | | | | | | |
| Disallowed (%) | | | | | | | | 0.11 | | | | | | | |

| EMDB | Unfocused | LSU | SSU |
|---|---|---|---|
| | Full with tRNA (PDB 9EVS) | | |
| EMDB | EMD-50011 | EMD-50012 | EMD-50013 |
| **Data collection and processing** | | | |
| Magnification | | ????x | |
| Voltage (kV) | | 300 | |
| Electron exposure (e–/Å$^2$) | | 46 | |
| Defocus range (μm) | | -0.5 to -2.0 | |
| Pixel size (Å) | | 1.058 | |
| Symmetry imposed | | C1 | |
| Initial particle images (no.) | | 141,939 | |
| Final particle images (no.) | | 83,443 | |
| Map resolution (Å) - FSC 0.143 | 3.05 | 2.95 | 3.08 |
| Map resolution range (Å) | 2.6 - 10.3* | 2.6 - 10.3* | 2.7 - 10.5* |
| **Refinement** | | | |
| Initial model used (PDB code) | | AF2 - 6XYW | |
| CC Model vs. Data (mask) | 0.81 | | |
| Map sharpening *B* factor (Å$^2$) | -36.5 | -35.0 | -37.1 |
| Model composition | | | |
| Non-hydrogen atoms | | 211459 | |
| Residues: Protein - Nucleotide | | 13740 - 4713 | |
| Water | | - | |
| Ligands | | ZN: 2<br>K: 74<br>MG: 357<br>ATP: 1<br>CLM: 1 | |
| *B* factors (Å$^2$) | | | |
| Protein | | 0.00/283.77/67.44 | |
| Nucleotide | | 1.69/341.90/82.12 | |
| Ligand | | 3.78/141.34/41.08 | |
| R.m.s. deviations | | | |
| Bond lengths (Å) | | 0.007 | |
| Bond angles (°) | | 0.989 | |
| Validation | | | |
| MolProbity score | | 1.39 | |
| Clashscore | | 3.53 | |
| Poor rotamers (%) | | 0.03 | |
| Ramachandran plot | | | |
| Favored (%) | | 96.30 | |
| Allowed (%) | | 3.61 | |
| Disallowed (%) | | 0.09 | |

| EMDB | Unfocused | SSU head | SSU body | RsgA |
|---|---|---|---|---|
| | SSU with RsgA (PDB 9EVT) | | | |
| EMDB | EMD-50014 | EMD-50015 | EMD-50017 | EMD-50016 |
| **Data collection and processing** | | | | |
| Magnification | | 165,000x | | |
| Voltage (kV) | | 300 | | |
| Electron exposure (e–/Å$^2$) | | 50.96 | | |
| Defocus range (μm) | | -0.5 to -2.5 | | |
| Pixel size (Å) | | 0.729 | | |
| Symmetry imposed | | C1 | | |
| Initial particle images (no.) | | 1,896,051 | | |
| Final particle images (no.) | | 208,855 | | |
| Map resolution (Å) - FSC 0.143 | 2.27 | 2.06 | 2.17 | 2.07 |
| Map resolution range (Å) | 1.6 - 7.4* | 1.6 - 7.2* | 1.6 - 7.4* | 1.6 - 5.4* |
| **Refinement** | | | | |
| Initial model used (PDB code) | | AF2 - 6XYW | | |
| CC Model vs. Data (mask) | 0.8 | | | |
| Map sharpening *B* factor (Å$^2$) | -39.8 | -31.7 | -35.8 | -37.7 |
| Model composition | | | | |
| Non-hydrogen atoms | | 85774 | | |
| Residues: Protein - Nucleotide | | 6462 - 1560 | | |
| Water | | - | | |
| Ligands | | ZN: 1<br>K: 10<br>MG: 85<br>ATP: 1<br>GTP: 1 | | |
| *B* factors (Å$^2$) | | | | |
| Protein | | 0.00/274.51/54.34 | | |
| Nucleotide | | 0.00/226.44/51.37 | | |
| Ligand | | 12.44/76.98/39.22 | | |
| R.m.s. deviations | | | | |
| Bond lengths (Å) | | 0.004 | | |
| Bond angles (°) | | 0.966 | | |
| Validation | | | | |
| MolProbity score | | 1.33 | | |
| Clashscore | | 3.86 | | |
| Poor rotamers (%) | | 0.02 | | |
| Ramachandran plot | | | | |
| Favored (%) | | 97.14 | | |
| Allowed (%) | | 2.81 | | |
| Disallowed (%) | | 0.05 | | |

* min to 75th in cryoSPARC Local Resolution Estimation (FSC 0.5)

### Supplementary Table 1 :
