## Supplementary Table 2 for "Structural insights into maturation and translation of a plant mitoribosome"

| Name of the protein | Arabidopsis UNIPROT | TAIR ID Arabidopsis | Modeled protein | ChainID | Location |
|---|---|---|---|---|---|
| uL1m | Q8RWT4 | At2g42710 | not modeled but visible | A | L1 stalk |
| uL2m C-ter | Q8VZU4 | At2g44065 | A0A0D3B6N5 B. oleracea var. oleracea | B | |
| uL2m N-ter mito | P93311 | AtMg00560 | A. thaliana | C | |
| uL3m | Q9LRN8 | At3g17465 | A0A0D3ACV4 B. oleracea var. oleracea | D | |
| uL4m | Q8VY61 | At2g20060 | A0A0D2ZRI2 B. oleracea var. oleracea | E | |
| uL5m | P42793 | AtMg00210 | A. thaliana | F | Central Protuberance |
| uL6m | Q9ZPX2 | At2g18400 | A0A0D3D316 B. oleracea var. oleracea | G | |
| bL9m | Q9LVU5 | At5g53070 | A. thaliana | H | |
| uL10m | Q9LHH1 | At3g12370 | A. thaliana | I | L7/L12 stalk |
| uL11m | Q9SVW7 | At4g35490 | A. thaliana | J | L7/L12 stalk |
| uL13m | Q7XA68 | At3g01790 | A0A0D3B978 B. oleracea var. oleracea | K | |
| uL14m | Q93Z17 | At5g46160 | A0A0D3D9E7 B. oleracea var. oleracea | L | |
| uL15m | Q9FLF3 | At5g64670 | A0A0D3BF13 B. oleracea var. oleracea | M | |
| uL16m | Q95747 | AtMg00080 | A. thaliana | N | |
| bL17m | Q8LD39 | At5g09770 | A0A8S9IRX0 Brassica cretica | O | |
| uL18m | Q8LDK5 | At5g27820 | A0A0D3DCH7 B. oleracea var. oleracea | P | Central Protuberance |
| bL19m | Q9T0D0 | At4g11630 | A0A0D3B0P2 B. oleracea var. oleracea | Q | |
| bL20m | Q8LCN1 | At1g16740 | A. thaliana | R | |
| bL21m | Q8L9A0 | At4g30930 | A. thaliana | S | |
| uL22m | Q8LDU0 | At4g28355 | A0A0D3BFS0 B. oleracea var. oleracea | T | |
| uL23m | Q9SMR5 | At4g39880 | A0A0D3A116 B. oleracea var. oleracea | U | |
| uL24m | Q9LT09 | At5g23535 | A. thaliana | V | |
| bL25-2m | Q9FKZ4 | At5g66860 | A. thaliana | W | Remodeled Domain III |
| bL25m | Q9SUR4 | At4g23620 | A0A0D3BK40 B. oleracea var. oleracea | X | |
| bL27m | Q94AC6 | At5g15220 | A0A0D3B184 B. oleracea var. oleracea | Y | Central Protuberance |
| bL28m | Q9SV23 | At4g31460 | A. thaliana | Z | |
| uL29m | Q94JQ7 | At1g07830 | A. thaliana | a | |
| uL30m | Q8L908 | At5g55140 | A. thaliana | b | |
| bL31m | Q1G3G5 | At5g55125 | A. thaliana | c | Central Protuberance |
| bL32m | Q944L5 | At1g26740 | A. thaliana | d | |
| bL33m | Q9SQT5 | At3g06320 | A. thaliana | e | Central Protuberance |
| bL34m | Q84R26 | At3g13882 | A0A0D3CKG1 B. oleracea var. oleracea | f | |
| bL35m | Q8LAA7 | At5g45590 | A. thaliana | g | |
| bL36m | Q8W464 | At5g20180 | A. thaliana | h | |
| mL40 | Q9M0V5 | At4g05400 | A0A3P6DV95 B. oleracea var. oleracea | i | Central Protuberance |
| mL41 | Q9LUJ9 | At5g40080 | A. thaliana | j | |
| mL43 | Q9M1A3 | At3g59650 | A0A0D3DUE5 B. oleracea var. oleracea | k | |
| mL46 | Q8L7U3 | At1g14620 | A. thaliana | l | Central Protuberance |
| mL53 | Q940G3 | At5g39600 | A. thaliana | m | L7/L12 stalk |
| mL54 | Q9S799 | At3g01740 | A. thaliana | o | L7/L12 stalk |
| mL59/mL64 | O65448 | At4g22000 | A. thaliana | p | Central Protuberance |
| mL60 | Q94F28 | At1g27435 | A0A0D3BIE3 B. oleracea var. oleracea | q | |
| mL80 | Q9C9B5 | At1g73940 | A0A0D3CWS6 B. oleracea var. oleracea | r | |
| mL87 | Q9SD44 | At3g51010 | A0A0D3DDW7 B. oleracea var. oleracea | s | |
| mL101 (rPPR4) | O22714 | At1g60770 | A0A0D3E593 B. oleracea var. oleracea | t | Remodeled Domain III |
| mL102 (rPPR5) | Q9ZUU3 | At2g37230 | A. thaliana | u | |
| mL104 (rPPR9) | Q9FME4 | At5g60960 | A. thaliana | v | Remodeled Domain III |

**Table 2 :** List of the LSU r-proteins

List of the r-proteins found in the LSU. Proteins are colored by conservation with the bacterial ribosome (blue) other mitoribosomes (yellow) or specific to the plant mitoribosome (red). Protein modeled in the density are indicated.
