## Supplementary Table 3 for "Structural insights into maturation and translation of a plant mitoribosome"

| Name of the protein | Arabidopsis UNIPROT | TAIR ID Arabidopsis | Modeled protein | ChainID | Location |
|---|---|---|---|---|---|
| uS2m | Q9GCB9 | At3g03600 | A0A0D3CN01 B. oleracea var. oleracea | A | body |
| uS3m | Q95749 | AtMg00090 | A. thaliana | B | head |
| uS4m | Q31708 | AtMg00290 | A. thaliana | C | body |
| uS5m | Q6GKU7 | At1g64880 | A0A0D3AS28 B. oleracea var. oleracea | D | body |
| bS6m | Q9LSA1 | At3g18760 | A0A0D3CIR5 B. oleracea var. oleracea | E | body |
| uS7m | P92557 | AtMg01270 | A. thaliana | F | head |
| uS8m | Q9M0E0 | At4g29430 | A. thaliana | G | body |
| uS9m | Q8L6Z4 | At3g49080 | A0A0D3DRJ4 B. oleracea var. oleracea | H | head and body |
| uS10m | P42797 | At3g22300 | A0A0D3CHA1 B. oleracea var. oleracea | I | head |
| uS11m | Q8VZT8 | At1g31817 | A. thaliana | J | body |
| uS12m | P92532 | AtMg00980 | A. thaliana | K | body |
| uS13m | Q9CA19 | At1g77750 | A0A0D3D1F0 B. oleracea var. oleracea | L | head |
| uS14m | Q9SMX4 | At2g34520 | A. thaliana | M | head |
| uS15m | Q9M8M9 | At1g80620 | A0A0D3D1Z7 B. oleracea var. oleracea | N | body |
| bS16m | Q9LTS6 | At5g56940 | A0A0D3ALM2 B. oleracea var. oleracea | O | body |
| uS17m | Q9LHN1 | At3g18880 | A. thaliana | P | body |
| bS18m | Q9LML3 | At1g07210 | A. thaliana | Q | body |
| uS19m | P39697 | At5g47320 | A. thaliana | R | head |
| bS21m | F4JCI2 | At3g26360 | A0A0D3AW14 B. oleracea var. oleracea | S | body |
| bTHXm | Q9SJU8 | At2g21290 | A0A0D3C0Z4 B. oleracea var. oleracea | T | head |
| mS23 | F4HPB7 | At1g26750 | A. thaliana | U | body |
| mS26 | Q945P2 | At5g49210 | A0A0D3DC57 B. oleracea var. oleracea | V | body |
| mS29 | Q8W4K2 | At1g16870 | A0A0D3CAV8 B. oleracea var. oleracea | W | head |
| mS31/mS46 | Q9C8L9 | At1g53645 | A. thaliana | X | head |
| mS33 | Q8GXH5 | At5g44710 | A. thaliana | Y | head |
| mS34 | Q9FHC3 | At5g52370 | A. thaliana | Z | body |
| mS35 | Q9LJQ6 | At3g18240 | A. thaliana | a | head |
| mS37 | A8MRX3 | At1g47278 | A. thaliana | b | head |
| mS38 | Q9FMK8 | At5g63150 | A. thaliana | c | body |
| mS41 | Q6IDB8 | At5g26800 | A0A0D3DCJ8 B. oleracea var. oleracea | d | body |
| mS45 | Q9LVA9 | At5g62270 | A0A0D3BFU2 B. oleracea var. oleracea | e | body |
| mS47 | Q8RXN4 | At4g31810 | A. thaliana | f | body |
| mS80 (rPPR6) | P0C896 | At3g02650 | A. thaliana | l | head extension |
| mS83 (rPPR10) | Q1JPP0 | At4g15640 | A. thaliana | h | foot |
| mS77 (NFD5) | A0A654ECT9 | At1g19520 | A. thaliana | i | body |
| mS76 (rPPR1) | Q8LE47 | At1g61870 | A. thaliana | j | foot |
| mS86 | Q9FZ84 | At1g18630 | A. thaliana | k | body |
| mS81 (rPPR8) | Q8LPF1 | At5g15980 | A. thaliana | g | head extension |
| RsgA | Q4V399 | At1g67440 | A. thaliana | 7 | |

**Table 3 :** List of the SSU r-proteins

List of the r-proteins found in the SSU. Proteins are colored by conservation with the bacterial ribosome (blue) other mitoribosomes (yellow) or specific to the plant mitoribosome (red). Protein modeled in the density are indicated.
