## Supplementary Table 4 for "Structural insights into maturation and translation of a plant mitoribosome"

| | Bacteria | Fl. Plants | | | Helix | Domain | Role |
|---|---|---|---|---|---|---|---|
| | Residue | Residue (Bo numbering) | Structure | Nanopore | | | |
| SSU | | A521 unknown | NO | YES | h18 | 5' | |
| | $m^{7}$G527 | **$m^{7}$G525** | YES | YES | h18 | 5' | decoding center |
| | | G527 unknown | NO | YES | h18 | 5' | |
| | $m^{2}$G966 | A959 unknown | NO | YES | h31 | 3'M | |
| | | A961 unknown | NO | YES | h31 | 3'M | |
| | $m^{4}$Cm1402 | **$m^{4}$Cm1664** | YES | NO | h44 (base) | 3'm | interacts with mRNA |
| | | U1804? | NO | YES | h44 (base) | 3'm | |
| | $m^{3}$U1498 | **$m^{3}$U1807** | YES | YES | h44 (h44-h45 linker) | 3'm | shapes mRNA channel |
| | $m^{2}$G1516 | **$m^{2}$G1825** | YES | YES | h45 | 3'm | supports P-stite conformation |
| | $m^{6}_{2}$A1518 | **$m^{6}_{2}$A1827** | YES | YES | h45 | 3'm | supports P-stite conformation |
| | $m^{6}_{2}$A1519 | **$m^{6}_{2}$A1828** | YES | YES | h45 | 3'm | supports P-stite conformation |
| LSU | Ψ746 | **Ψ891** | YES | NO | h35 | II | |
| | Ψ955 | **Ψ1105** | YES | NO | h39 | II | |
| | | **Ψ2188** | YES | YES | h68 | IV | |
| | Ψ1911 | Ψ2229 | N/A | YES | h69 | IV | |
| | $m^{3}$Ψ1915 | Ψ2233 | N/A | YES | h69 | IV | extends stacking and faces h44 of the small subunit |
| | Ψ1917 | Ψ2235 | N/A | YES | h69 | IV | |
| | | Ψ2239 | N/A | YES | h69 | IV | |
| | $m^{5}$U1939 | **$m^{5}$U2257** | YES | NO | h70 | IV | |
| | $m^{5}$C1962 | **$m^{5}$C2280** | YES | NO | h71 | IV | Extends G1945-C1972 stacking within the H70-H71 helical junction |
| | Gm2251 | **Gm2560** | YES | YES | h80 | V | P-loop, G2560 is a key residue which base- pairs with the CCA-end of the P-site tRNA |
| | D2449 | **$H_{2}$U2754** | YES | YES | ssRNA prior to h89 | V | |
| | Ψ2457 | **Ψ2762** | YES | NO | h89 | V | |
| | Cm2498 | **Cm2803** | YES | NO | h89 | V | |
| | $s^{2}$C2501 | $s^{2}$C2806 | NO | YES | ssRNA prior to h90 | V | |
| | $m^{2}$A2503 | **$m^{2}$A2808** | YES | YES | ssRNA prior to h90 | V | Peptide tunnel |
| | Ψ2504 | Ψ2809 | NO | YES | ssRNA prior to h90 | V | |
| | Um2552 | **Um2857** | YES | YES | h92 | V | A-loop, Intercalates between the bases of U2857 and G2858 (G2858 is a key residue which base-pairs with the CCA-end of the A-site tRNA) |
| | | Gm2858 | NO | YES | h92 | V | A-loop, G2858 is a key residue which base-pairs with the CCA-end of the A-site tRNA |
| | Ψ2580 | **Ψ2885** | YES | YES | h90 | V | stabilizes A-site tRNA |
| | Ψ2605 | **Ψ2910** | YES | YES | h93 | V | |

**Table 4 :** List of the rRNA modifications

Summary of the rRNA modifications detected in B. oleracea mitoribosome. B. oleracea contains 11 modified residues in the SSU and 20 in the LSU. The method of detection (Cryo-EM structure or Nanopore DRS), their helix and domain position on the rRNA, as well as their functional role are presented for each modified residue.
