## Supplementary Table 5 for "Structural insights into maturation and translation of a plant mitoribosome"

| Oligonucleotide | Sequence |
|---|---|
| 26S_F | CGAAAAGAATGCATTGGAT |
| 26S_R | TCGTTTAGTACGAGATGGC |
| 18S_F | ATCATAGTCAAAAGAAGAGTTTG |
| 18S_R | GGATTCAATCCAGCCACAG |
| T7-26S | TAATACGACTCACTATAGCGAAAAGAATGCATTGGAT |
| T7-18S | TAATACGACTCACTATAGATCATAGTCAAAAGAAGAGTTTG |
| RTA_OligoA_BC1 | /5PHOS/GGCTTCTTCTTGCTCTTAGGTAGTAGGTTC |
| RTA_OligoB_26S_BC1 | GAGGCGAGCGGTCAATTTTCCTAAGAGCAAGAAGAAGCCTCGTTTAGTA |
| RTA_OligoB_18S_BC1 | GAGGCGAGCGGTCAATTTTCCTAAGAGCAAGAAGAAGCCGGATTCAATC |
| RTA_OligoA_BC3 | /5PHOS/GTACTTTTCTCTTTGCGCGGTAGTAGGTTC |
| RTA_OligoB_26S_BC3 | GAGGCGAGCGGTCAATTTTCCGCGCAAAGAGAAAAGTACTCGTTTAGTA |
| RTA_OligoA_BC2 | /5PHOS/GTGATTCTCGTCTTTCTGCGTAGTAGGTTC |
| RTA_OligoB_18S_BC2 | GAGGCGAGCGGTCAATTTTCGCAGAAAGACGAGAATCACGGATTCAATC |
| Barcode 01 (BC1) | GGCTTCTTCTTGCTCTTAGG |
| Barcode 02 (BC2) | GTGATTCTCGTCTTTCTGCG |
| Barcode 03 (BC3) | GTACTTTTCTCTTTGCGCGG |

**Supplementary Table 5 :** List of oligonucleotides

| FLO-PRO002 run | Barcode ID | Target rRNA | direct RNA-seq reads available |
|---|---|---|---|
| TGS104_16052024 | Barcode01 | Native 26S | 17 849 |
| TGS104_14052024 | Barcode03 | IVT 26S | 680 928 |
| TGS103_21032024 | Barcode01 | Native 18S | 2 463 135 |
| TGS103_21032024 | Barcode02 | IVT 18S | 1 587 154 |
| **FLO-PRO002 run** | **Barode ID** | **Target rRNA** | **Selected full-length reads** |
| TGS104_16052024 | Barcode01 | Native 26S | 2166 |
| TGS104_14052024 | Barcode03 | IVT 26S | 2190 |
| TGS103_21032024 | Barcode01 | Native 18S | 2000 |
| TGS103_21032024 | Barcode02 | IVT 18S | 2000 |

**Supplementary Table 6 :** Read statistics from Deeplexicon demultiplexing, length filtering and read selection
